# Consequences of coupled barriers to gene flow for the build-up of genomic differentiation

**DOI:** 10.1101/2021.09.02.458401

**Authors:** Henry D. Kunerth, Steven M. Bogdanowicz, Jeremy B. Searle, Richard G. Harrison, Brad S. Coates, Genevieve M. Kozak, Erik B. Dopman

**Affiliations:** Department of Ecology and Evolutionary Biology, Cornell University; 02747USDA-ARS, Corn Insects & Crop Genetics Research Unit, Ames, IA, 50011; Department of Biology, UMass-Dartmouth, Dartmouth, MA; Department of Biology, Tufts University, Medford, MA, 02155

**Author notes:** Corresponding author: Henry D. Kunerth, A406B Corson Hall, 215 Tower Road, Ithaca NY, 14853. Kunerth, Harrison, Kozak, and Dopman conceived of the project. Kunerth, Kozak, Coates and Bogdanowicz contributed to sample preparation. Kunerth and Bogdanowicz performed PCR and sequencing library preparation of targeted marker dataset. Kunerth and Dopman performed data analysis on the targeted marker dataset. Kozak and Coates performed pool-seq library preparation, and Kozak and Dopman analyzed the pooled sequencing dataset. Kunerth, Searle, Kozak, and Dopman wrote the first draft of the manuscript and Kunerth and Dopman revised the manuscript with contributions from other authors. We are grateful to the substantial contributions of many lab members and colleagues for their comments, criticisms, and discussions. Jessie Martin, Crista Wadsworth, Gabriel Golczer, Benjamin Hamilton, and Charles Mason all contributed to field collections, Christopher Watson assisted in the lab, and Qi Sun helped with bioinformatic analysis and scripting. H.D.K received Fellowship support from the Presidential Life Sciences and Cornell Fellowships. Partial research support was provided by USDA, ARS CRIS Project 5030-22000-019-00D. Any mention of products or services is for research purposes only, and does not constitute an endorsement for use by the USDA. USDA is an equal opportunity provider and Employer. E.B.D., R.G.H., and J.B.S. were funded by the National Science Foundation, and E.B.D. by Tufts University and cooperative agreement 58-5030-7-066 between USDA-ARS and Tufts University. All data is archived in GenBank: PRJNA361472; PRJNA540655; PRJNA655940; PRJNA656263; PRJNA794858.

## Abstract

Theory predicts that when different barriers to gene flow become coincident, their joint effects enhance reproductive isolation and genomic divergence beyond their individual effects, but empirical tests of this ‘coupling’ hypothesis are rare. Here, we analyze patterns of gene exchange among populations of European corn borer moths that vary in the number of acting barriers, allowing for comparisons of genomic variation when barrier traits or loci are in coincident or independent states. We find that divergence is mainly restricted to barrier loci when populations differ by a single barrier, whereas the coincidence of temporal and behavioral barriers is associated with divergence of two chromosomes harboring barrier loci. Furthermore, differentiation at temporal barrier loci increases in the presence of behavioral divergence and differentiation at behavioral barrier loci increases in the presence of temporal divergence. Our results demonstrate how the joint action of coincident barrier effects leads to levels of genomic differentiation that far exceed those of single barriers acting alone, consistent with theory arguing that coupling allows indirect selection to combine with direct selection and thereby lead to a stronger overall barrier to gene flow. Thus, the state of barriers – independent or coupled – strongly influences the accumulation of genomic differentiation.

## INTRODUCTION

Understanding the origin of new species is crucial for explaining patterns of biodiversity. New species are thought to arise as populations diverge and accumulate trait differences that limit interbreeding, genetic exchange, and sharing of evolutionary processes (Mayr 1942, Coyne and Orr 2004, Harrison and Larson 2014). Consequently, there is long-standing interest in understanding how barriers to reproduction evolve and promote independent evolution of populations in nature, as characterized by the emergence of distinct genetic groups.

Most sister species are separated by multiple barriers to gene flow such as differences in mating time, differences in mate choice, and decreased hybrid survival (Coyne and Orr 2004; Dopman et al. 2010), arguing that speciation depends on the accumulation of barriers and strong reproductive isolation. Different barriers need not, however, initially evolve between the same populations. Instead, independent barriers could restrict gene flow between different populations, potentially as a gradual build-up of barriers with geographic distance (Barton 2013). Without the coincident action of multiple barriers between the same populations, theory indicates that differentiation will be centered at loci underlying barrier traits, reflecting weak overall reproductive isolation and potentially migration-selection balance (Barton 1983; Barton and Bengtsson 1986; Barton and de Cara 2009; Butlin and Smadja 2018; Flaxman et al. 2014; Schilling et al. 2018). At other genomic regions, gene flow and recombination can allow for shared evolutionary processes within what is essentially a single population (Barton 2013).

When multiple barrier traits become coincident in a process known as ‘coupling,’ their joint effects are believed to enhance reproductive isolation and genomic divergence beyond their individual effects, potentially leading to strong reproductive isolation sufficient for the origin of new species (Barton 1983; Barton and Bengtsson 1986; reviewed in Butlin and Smadja 2018). Recent simulations of divergence with gene flow suggests the presence of tipping-point dynamics, wherein a certain threshold of coupling, achieved by the accumulation of new barrier loci or increased linkage disequilibrium (LD) among existing loci, initiates a feedback loop of rapid differentiation across the genome (Flaxman et al. 2014, Schilling et al. 2018, reviewed in Nosil et al. 2017). This coupling threshold may correspond to a critical point at which the strength of selection across accumulated barrier loci, arising from both direct fitness effects of each locus and indirect fitness effects of their non-random associations, outweighs the homogenizing impact of recombination (measured as *φ*, the ‘coupling coefficient’, Barton 1983). As coupling builds towards this tipping point, recombination is predicted to cause a lag in divergence at neutral loci resulting in heterogeneous genomic differentiation (Nosil et al. 2017; Schilling et al. 2018). Genomic architectures that reduce recombination, including pleiotropy, tight physical linkage, or chromosomal rearrangements, are therefore expected to promote coupling (Servedio et al. 2011; Dagilis and Kirkpatrick 2016, Ortiz-Barrientos et al. 2016; Kirkpatrick & Barton 2006). Although coupling does not require physical linkage, in a widespread pattern known as the ‘large X-effect,’ sex chromosomes often harbor more barrier loci than autosomes (Coyne and Orr, 1989; Presgraves 2008; 2018). Consequently, the X (or Z) chromosome might be prone to coupling dynamics and rapid chromosomal differentiation (Lasne et al. 2017; Presgraves 2018). Consistent with this, 95% of 129 studies reviewed in Presgraves (2018) find evidence between taxa of elevated differentiation on sex chromosomes relative to autosomes, although there is no shortage of alternative hypotheses for biased sex chromosome differentiation (e.g. Caballero 1995; Pool & Nielsen 2007; Charlesworth 2012; reviewed in Presgraves 2018).

Critical to theoretical predictions of a build-up of genomic divergence that ultimately characterizes new species is the coupling of multiple barriers to gene flow. Understanding how coupling counteracts gene flow to allow genomic divergence requires us to reconstruct the sequence of events, from barrier independence to barrier coincidence, as speciation progresses. Much empirical work has indirectly studied the evolution of coupling during speciation by quantifying genomic differentiation across lineages at differing stages of divergence (Gagnaire et al. 2013, Seehausen et al. 2014; Shaw & Mullen 2014; Riesch et al. 2017; Xu & Shaw 2019, 2021). Equally valuable are systems allowing direct comparisons of genomic variation when barrier traits or loci are found in either a coincident or independent state. Such direct studies of coupling dynamics may be especially challenging in many systems because barriers to gene flow, trait loci, and recombination landscapes are either unknown or barriers are found exclusively in a coincident state. Consequently, strong empirical tests of coupling dynamics are rare.

The European corn borer moth (ECB) *Ostrinia nubilalis* (Hübner) (Lepidoptera: Crambidae) is useful for studying coupling dynamics because hybridizing population pairs can differ by independent or coincident barriers. In addition to non-coincident barrier effects (Butlin and Smadja 2017), ECB populations differ in combinations of sex pheromone, host-plant association, and phenology (Malausa et al., 2007; Dopman et al. 2010, Coates et al. 2018). Throughout much of its introduced range in North America and its ancestral range in Europe and Asia, two pheromone strains co-exist (Klun and Cooperators, 1975; Anglade and Stockel 1984). In the E strain, females produce and males respond to a sex pheromone comprised mainly of the E isomer of 11-tetradecenyl acetate (E pheromone), whereas Z moths use a blend of mainly Z isomer (Z pheromone) (Klun 1973, Kochansky 1975). The Z strain is more widespread, including areas west of the Appalachian Mountains where it occurs alone (Klun and Cooperators, 1975; Sorenson et al. 1992). Although dietary generalists (Hodgson, 1928; Lewis, 1975), larvae show some host-plant specialization on cultivated maize (Calcagno, et al. 2007; Fisher et al 2017), their predominant host (O’Rourke et al. 2010; Coates et al. 2019). However, in France, the E strain shows a near complete affiliation with non-maize hosts (Malausa et al., 2007), resulting in its identification as the sister species *O. scapulalis* (Calcagno et al. 2007; Frolov et al. 2007). Finally, three locally adaptive ecotypes have been described in North America that differ in seasonal phenology and voltinism. A northern ecotype (> 42°N) produces one mid-season generation per year (univoltine), a central ecotype (~36–45°N) makes one early-season and one late-season generation (bivoltine), and a southern ecotype (< 40°N) makes three or more generations per year (multivoltine) (Showers et al. 1976; 1981). The northern ecotype uses the Z pheromone, whereas both Z and E pheromones are used by central and southern ecotypes (Sorenson et al. 1992; Showers 1993). Northern and central ecotypes become sympatric across an area stretching from Wyoming to Maine (Showers et al. 1975; 1981), resulting in population pairs differing by voltinism alone, or both voltinism and pheromone.

The three main axes of phenotypic divergence (phenology, pheromone signaling, host-plant use) result in various barrier effects. Populations of North American ECB moths can experience a temporal barrier from differences in seasonal phenology (bivoltine versus univoltine), a behavioral barrier from differences in pheromone communication (Z versus E), or a coincidence of the two barrier types (bivoltine E versus univoltine Z) (Figure 1). Ten other potential barriers to gene flow have been studied that are comparatively weaker (Dopman et al. 2010). North American populations are derived from Hungary and Italy (Smith 1920, Caffrey and Worthly 1927) rather than France, where barrier effects between the ECB moth *O. nubilalis* and its sister species *O. scapulalis* consist of a coincidence of pheromone and host, but not voltinism (Thomas et al. 2003).

**Figure 1.**
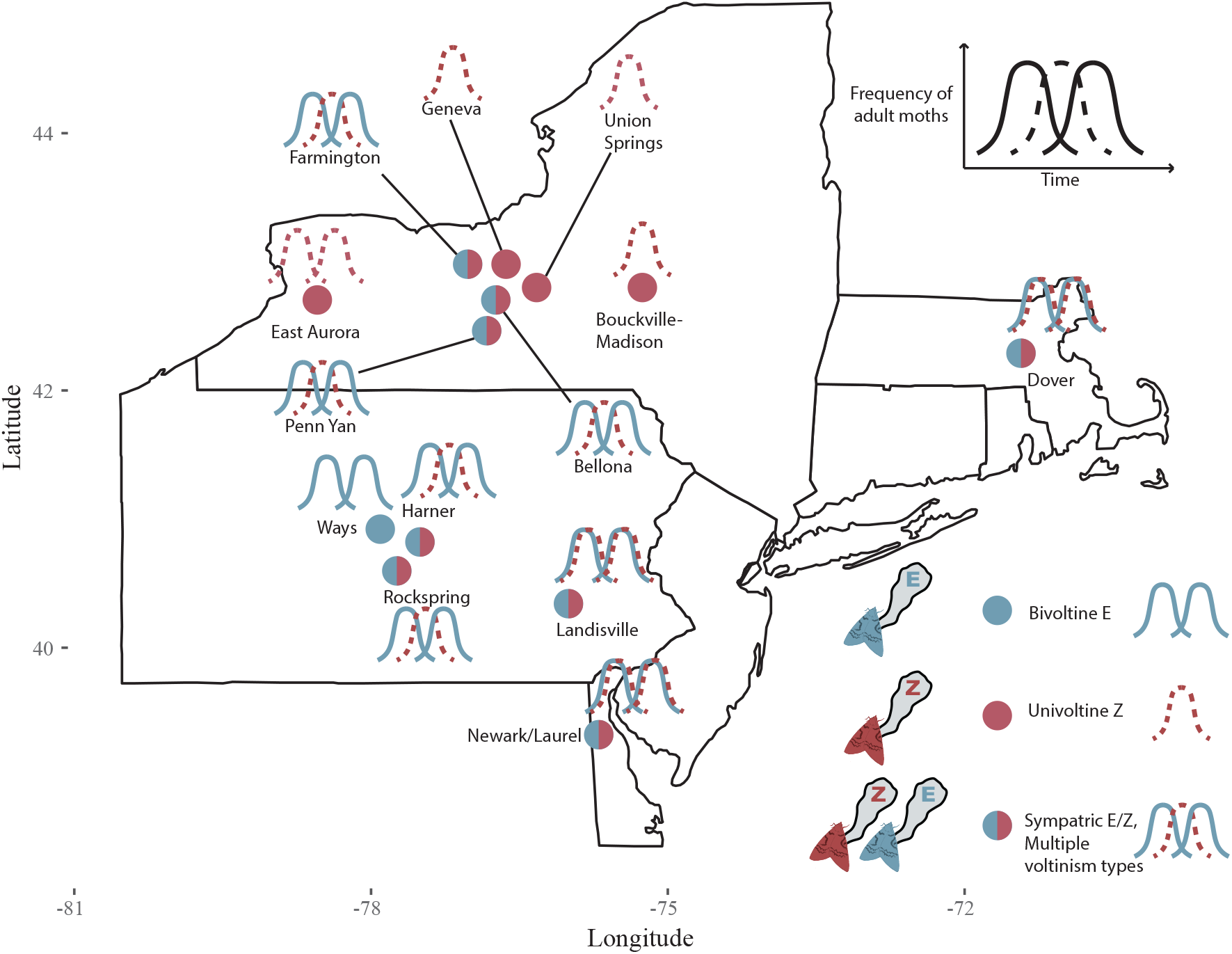
Map of collection sites and phenotypic information. Graphs indicate the frequency of adult moths across the growing season and number of distinct generations (voltinism). Graph and circle color indicates whether there were E (blue), Z (red) or both pheromone strains present at each collection site.

Four other aspects of the system are noteworthy because they have been shown by simulations to efficiently reduce gene flow or promote coupling (Flaxman et al. 2014, Nosil et al. 2017, Schilling et al. 2018). First, evidence suggests pheromone signal and preference, controlled by *pgFAR* (pheromone production) and *bab* (preference) genes (Unbehend et al. 2021, Lassance et al. 2010), are ‘multiple-effect traits’ (Smadja and Butlin 2011) that contribute to both pre- and post-zygotic isolation (mate choice and behavioral sterility) (Glover et al. 1991; Dopman et al. 2010; Unbehend et al. 2021). Second, in an example of ‘magic trait’ evolution (Servedio et al. 2011), temporal isolation arising from non-random meeting of univoltine and bivoltine adults is an automatic, pleiotropic outcome of divergent selection on diapause phenology, controlled by two interacting clock genes *per* and *Pdfr* (Dopman et al. 2005; Levy et al. 2015; Kozak et al. 2019). Third, three of four identified barrier loci (*per*, *Pdfr*, *bab*) map to the Z chromosome, creating a ‘large Z-effect’ (Dopman et al. 2004; Kozak et al. 2019; Unbehend et al. 2021). Finally, a chromosomal rearrangement (putatively an inversion(s)) suppressing recombination across ~40% of the Z chromosome has captured at least one barrier locus (*per*) and may be abundant within some E pheromone populations (Wadsworth et al. 2015; Kozak et al. 2017).

Prior population genetic results suggest identified barriers influence the build-up of genomic divergence. Geographic distance is weakly associated with genetic variation throughout the range (Bourguet et al. 2000; Kim et al. 2011; Coates et al. 2019), consistent with numerous examples of long-distance dispersal (30–60 km) (reviewed in Sappington 2018). Instead, genetic structure appears related to barrier effects. In France, differences in pheromone and host use between sympatric *O. nubilalis* and *O. scapulalis* are associated with significant genetic differentiation (Bethenod et al., 2005; Malausa et al., 2007). In an analysis restricted to 65 mostly autosomal SNPs among 12 localities in North America, pheromone signaling but not host or geography significantly contribute to genetic differentiation, but the influence of phenology was unmeasured (Coates et al. 2019). Kozak et al. (2017) used genome-wide data to study the Z chromosome rearrangement in North America and did find elevated differentiation when populations differed in both pheromone and phenology compared to pheromone alone, but contrasts were between single population pairs. Overall, these prior studies do imply that barrier phenotypes heavily shape patterns of gene flow. However, incomplete genomic data or barrier effect combinations have made it difficult the evaluate the importance of coupling dynamics in this system.

Here, we help resolve some earlier limitations by analyzing pooled genome data from 14 population samples and compare genetic divergence between four types of population pairs characterized by 1) no barrier effects, 2) only phenology differences (voltinism), 3) only pheromone differences, or 4) differences in both phenology and pheromone (joint divergence). In addition, we analyze targeted amplicon sequencing of the ‘large-effect’ Z chromosome and the autosomal pheromone production locus *pgFAR* from individuals collected from six locations varying in barrier number. Critically, these data allow for comparisons of sympatric populations differing by independent or coincident barriers, allowing us to ask whether we can detect empirical evidence that coupling promotes speciation as suggested by theory. Accordingly, compared to barrier traits when acting alone, upon coincidence we predict evidence of stronger selection or more efficient reduction of gene flow. Two expected signatures are increased divergence around individual barrier loci as well as enhanced coupling (linkage disequilibrium, LD) among barrier loci. If overall selection after coupling outweighs recombination, theory predicts tipping point dynamics and a transition towards differentiation of neutral loci. Differentiation can be genome-wide or more localized, but the combination of a ‘large Z-effect’ and a Z-linked rearrangement might suffice in allowing for the build-up of sex chromosome divergence. An alternative possibility is that coincident barriers do not lead to a detectable strengthening of the overall barrier. This could mean that individual barriers are so weak that they evolve independently even when coincident, or possibly that coupling effects are difficult to detect in nature, and theory could explore further quantitative predictions for empirical systems.

## MATERIALS AND METHODS

We sampled 13 localities in Eastern United States between 2000 and 2014 (Figure 1). Pheromone usage was based on male capture in traps baited with either synthetic E (“New York”) or Z (“Iowa”) pheromone lures (*Heliothis* traps, Scentry Biologicals, Billings, MO). Traps were separated by ≥ 12 m next to sweetcorn fields. Trap data were verified by gas chromatography of female pheromone gland extracts for the presence of E and Z isomers. For 7 sites, we sampled diapausing larvae from corn stalks and raised individuals under diapause breaking conditions (26°C, 16h light) until eclosion, allowing classification of voltinism based on post-diapause development (PDD) time (bivoltine: PDD time < 18 days; univoltine: PDD time > 38 days). For one population (Dover), we collected direct developing larvae in July. These individuals were mated as adults, larvae were exposed to diapausing inducing conditions and then phenotyped for PDD. For 5 sites, we determined voltinism by the seasonal timing of male capture in pheromone traps (bivoltine = early and late flight; univoltine = single mid-summer flight) (Glover 1991, Calvin 1994, Zaman 2008, Dopman 2010).

### Pooled genome sequencing

To evaluate how the transition between independent and joint axes of phenotypic divergence might affect genomic differentiation, we performed pooled genome sequencing on 14 population samples (425 individuals) separated by pheromone and phenology (5 bivoltine E (BE) populations, 3 bivoltine Z (BZ) populations, and 6 univoltine Z (UZ) populations; see Table 1). Pheromone strain assignment used *pgFAR* genotype (E or Z homozygotes) from individuals as determined by a diagnostic restriction enzyme digest of *pgFAR* PCR products (Coates et al. 2013). DNA was isolated using Qiagen DNeasy tissue kits (Qiagen, Germantown, MD) without vortexing to preserve high molecular weights, and samples were treated with RNase A (Qiagen). DNA from samples from Bellona, NY were extracted using Qiagen genomic tips (20 G). After quantifying DNA concentration using a Qubit (Thermo-Fisher), pools combined equal amounts of DNA per individual (Table 2). Pooled libraries (2 libraries per lane; 2-5 μg/uL per pool) were sequenced on an Illumina HiSeq3000 (2×150 bp) (except for Rockspring and Ways, which had 1×100 bp data) at Iowa State University DNA Facility, Ames, IA.

**Table 1.** Characteristics and locations for pooled data.

| Site | Pheromone | Voltinism | Number of males | Number of females | Collection method | Coverage after alignment |
| --- | --- | --- | --- | --- | --- | --- |
| Bouckville-Madison, NY | Z | Univoltine | 14 | 6 | DI | 37.0 |
| Bellona, NY | E | Bivoltine | 31 | 0 | PT | 53.18 |
| Bellona, NY | Z | Univoltine | 25 | 0 | PT | 41.52 |
| Dover, MA | Z | Bivoltine | 17 | 5 | DD | 15.83 |
| East Aurora, NY | Z | Bivoltine | 16 | 18 | DI | 32.35 |
| Geneva, NY | Z | Univoltine | 13 | 12 | DI | 5.94 |
| Harner, PA | E | Bivoltine | 31 | 0 | PT | 7.25 |
| Harner, PA | Z | Univoltine | 27 | 0 | PT | 4.59 |
| Landisville, PA | E | Bivoltine | 41 | 0 | PT | 19.29 |
| Landisville, PA | Z | Bivoltine | 39 | 0 | PT | 5.04 |
| Penn Yan, NY | Z | Univoltine | 11 | 15 | DI | 23.54 |
| Rockspring, PA | E | Bivoltine | 34 | 0 | PT | 10.36 |
| Rockspring, PA | Z | Univoltine | 33 | 0 | PT | 10.26 |
| Ways, PA | E | Bivoltine | 37 | 0 | PT | 13.33 |

PT = pheromone trap, DD = direct-developing larvae, DI = Diapausing larvae

**Table 2.** Characteristics and locations for amplicon sequencing data.

| Site | Population Type | Number of females | Number of males | Frequency of *pgFAR* genotypes |  |  | Difference in average eclosion time between *pgFAR* ZZ and EE insects (days) <sup>†</sup> |
| --- | --- | --- | --- | --- | --- | --- | --- |
|  |  |  |  | EE | EZ | ZZ |  |
| Dover, MA | Bivoltine E / Bivoltine Z | 22 | 27 | 50.0% | 29.5% | 20.5% | ‡ |
| East Aurora, NY | Bivoltine Z | 18 | 15 | 0.0% | 3.5% | 96.5% | - |
| Farmington, NY | Bivoltine E / Univoltine Z | 26 | 19 | 27.8% | 8.3% | 63.9% | 20.10 |
| Laurel & Newark, DE | Bivoltine E / Bivoltine Z | 51 | 0 | 12.8% | 34.0% | 53.2% | 1.74 |
| Penn Yan, NY | Bivoltine E / Univoltine Z | 20 | 9 | 24.0% | 16.0% | 60.0% | 34.63 |
| Union Springs, NY | Univoltine Z | 24 | 29 | 0.0% | 0.0% | 100.0% | - |

<sup>†</sup> Time to adult eclosion for diapausing insects (26°C, 16h light) acts as a proxy for post-diapause development time
<sup>‡</sup> Time to eclosion data was not available for these samples, but was calculated for the subsequent F<sub>1</sub> generation in the lab and confirmed to be bivoltine (see Material and Methods for more detail).

Trimmomatic v.35 removed Illumina adapters (TruSeq2 single-end or TruSeq3 paired-end), and reads with quality score (*q*) < 15 over a sliding window of 4 and reads < 36 bp long were removed (Bolger et al. 2014). Repetitive regions of the ECB reference genome (GenBank BioProject: PRJNA534504; Accession SWFO00000000; Kozak et al. 2019) were masked by RepeatMasker (using *Drosophila melanogaster* TE library in repbase; http://www.repeatmasker.org/; accessed March 2017), to which trimmed genomic data were aligned using bowtie2 v.2.2.3 (Langmead and Salzberg 2012). Aligned reads were sorted and filtered using Picard (v.2.8.0; http://broadinstitute.github.io/picard/) and SAMtools (v.0.1.18) to remove duplicates and reads with a mapping quality score (*Q*) < 20 (Li et al. 2009). We calculated coverage from the aligned, filtered, sorted bam files using bedtools v.2.26.0 and the genomeCoverageBed function (Table 2) (Quinlan 2014). As described in Kozak et al. (2019), we determined chromosomal positions of scaffolds by aligning genotyping-by-sequencing data from pedigree families (with and without the recombination suppressor segregating) to the genome scaffolds (Coates and Siegfried 2015, Kozak et al. 2017) and constructing linkage groups from identified SNPs in R v.3.5.1 (R Core Team, 2018) using the R/qtl package (Broman et al. 2003). We considered SNPs from Z chromosome-linked scaffolds that showed no recombination in E x Z strain pedigrees but recombined in Z x Z strain pedigrees to be inside the region of suppression (Wadsworth et al. 2015; Kozak et al. 2017). Twenty-nine scaffolds comprising 8.96 Mb of sequence were determined to be inside the region of recombination suppression while 38 scaffolds (11.94 Mb) were outside the region. We placed a small number (3%) of the Z chromosome-linked scaffolds using positions relative to other markers if they were identified as best hits from BLASTN searches using BLAST v.2.8.1 against the genome scaffolds of markers or BACs from previously published Z-linkage maps and pedigree families in *O. nubilalis* (Kroemer et al. 2009; Levy et al. 2015; Koutroumpa et al. 2016; Yasukochi et al. 2016) and also in BLASTN searches of Z chromosome-linked genome scaffolds in the sister species *O. scapulalis* against the ECB genome (Brousseau et al. 2018). Order and orientation of scaffolds was determined using linkage information from previous maps and gene order from *Bombyx mori* and approximate locations across the Z chromosome was calculated in Mb (described in Kozak et al. 2019). Scaffolds with no evidence for Z chromosome linkage in any pedigree were designated as autosomal. Chromosome assignment of scaffolds to various autosomes (chr. 2–31) used linkage groups in the pedigree families, and synteny of scaffolds to *Bombyx mori* and *Melita cinxia* chromosomes (described in Kozak et al. 2019). Chromosome 12 contains the *pgFAR* gene (Dopman et al. 2004).

We used SAMtools to identify single nucleotide polymorphisms (SNPs) in populations (Li et al. 2009). Scripts from Popoolation2 v.1.201 were used to filter (removing SNPs near small indels, minor alleles that did not appear twice in each population), calculate allele frequency, and *F*_ST_ (Kofler et al. 2011). Pairwise differences in *F*_ST_ were calculated on 100 kb non-overlapping sliding windows (including only windows with a minimum coverage > 14 reads and a maximum coverage < 300 and at least 20 polymorphisms within the window, as recommended by Fabian et al. 2012). We conducted pairwise *F*_ST_ comparisons among four categories of populations: no divergence in pheromone or phenology (BZ vs. BZ, UZ vs. UZ), differing only in voltinism (BZ vs. UZ), differing only in pheromone (BE vs. BZ), and differing in both pheromone and voltinism (BE vs. UZ).

We separately compared *F_ST_* for autosomal and Z chromosomal loci, and Z loci located inside the region of suppressed recombination (“rearranged”) or Z loci located outside this region (“collinear”). We removed and separately analyzed genomic scaffolds containing loci known to contribute to voltinism [*per* (voltinism; scaffold 532; Z chromosome inside “rearranged” region; Kozak et al. 2019), and *Pdfr* (voltinism; scaffold87, Z chromosome “collinear” region; Kozak et al. 2019)] and pheromone trait differences [*pgFAR* (female pheromone production: scaffold178, chr 12; Lassance et al. 2009); *bab* (male flight response; scaffold662, scaffold50; Z chromosome “collinear” region; Unbehend et al. 2021)]. We calculated and plotted *F_ST_* separately for 100 kb windows overlapping seasonal timing genes (*per*, *Pdfr*, N = 4 windows total, 2 in each scaffold), windows overlapping the *bab* locus (N = 4 windows), and windows overlapping the *pgFAR* locus (N = 1 window). We calculated the average *F_ST_* for all population pairs of a given type within each 100 kb window and then took the mean of all windows for each chromosome or genomic region. We performed comparison among population types using a Kruskal-Wallis test and pairwise posthoc Dunn’s tests in the PMCMRplus package (v1.4.4) with a false discovery rate correction in R (Pohlert 2014).

As some population pairs were collected from the same site (“sympatric”) and some from geographically distant sites (“allopatric”), we tested for an effect of geographic distance on the matrix of all pairwise *F*_ST_ differences using Mantel tests in the R package vegan v.2.5.6 (Oksanen et al. 2019). We used a matrix of geographic distance between sites (circle distance) as the explanatory matrix. We also performed a conservative analysis that was restricted to those population pairs within ~100 km (1-3 generations of dispersal, Sappington 2018) (6 population samples in NY, 7 population samples in PA).

### Targeted amplicon sequencing

To specifically test how the transition from independent to coincident barrier effects in sympatry influences the build-up of genomic differentiation, we performed amplicon sequencing of the ‘large-effect’ Z chromosome and *pgFAR* on population samples at six locations (260 individuals, 6 sites) experiencing either no phenotypic divergence or barrier effects (1 allopatric BZ population, 1 allopatric UZ population), pheromone divergence and behavioral isolation (2 sympatric BE/BZ population pairs), or joint phenotypic divergence and cumulative temporal and behavioral isolation (2 sympatric BE/UZ population pairs) (Table 2). Females were classified by gas chromatography of pheromone gland extracts, and both sexes by diagnostic SNPs at the *pgFAR* gene. Phenology classification is described above, using adult eclosion time as a proxy for PDD (except at Dover as described above). At Delaware, only female samples were analyzed, but all other locations had both male and female samples in approximately equal proportion.

We used 67 primers for multiplex PCR of 450–470 bp product sizes (Table S1). Primers were distributed across the Z chromosome, including near one of the two temporal barrier genes (*Pdfr*), and one near *pgFAR* on chromosome 12 (Figure S1). Only loci successfully amplified from both pheromone strains were carried forward to the final multiplex. We carried out two multiplexed reactions using a Qiagen multiplex PCR kit in two 384-well plates. Following amplification, we pooled products and transferred to one 384-well plate for barcoding. We created a single library by pooling these products after dual barcoding using Nextera N5/N7 primers and OneTaq Polymerase (New England Biolabs). We then cleaned the library using Ampure XP (Beckman Coulter) for 350–600 bp product sizes. We quantified libraries using a Qubit and sequenced them on a single Illumina MiSeq (2×250 bp) lane at the Cornell Institute of Biotechnology Genomics Facility, Ithaca, NY.

We demultiplexed and trimmed raw reads using a quality threshold of 20 with a custom Perl script. For each locus, we aligned reads using ClustalW (Larkin et al. 2007), generated a haplotype table for each sample at all loci, and called genotypes from the two most common haplotypes (minimum 4 reads total). If the ratio of the second most common genotype to the most common genotype was below 0.2, a homozygous genotype for the most common allele was called. We removed from analysis all individuals with ≥ 50% missing data, then loci with > 35% missing data. Following these cutoffs, we re-filtered all individuals and removed those with ≥ 35% missing data. We tested lower missingness cutoffs but found that a 35% missingness filter produced STRUCTURE and PCA results with no discernable differences to lower filters. Applying quality filters resulted in 217 moths (85 males, 132 females).

Using Z chromosome data, we inferred the presence of distinct populations, assigned individuals to populations, and identified admixed individuals using STRUCTURE (v2.3.4) (Pritchard et al. 2000). Since hemizygous females only carry one Z chromosome, we treated all samples as haploid with males contributing two haploid sets. To assure that this design did not bias results, we conducted separate STRUCTURE runs using only male samples (treated as diploid) or only female samples (treated as haploid). In all cases, we demonstrated qualitatively similar results and could not detect any bias. The presented results include all samples and all markers, unless otherwise specified. We tested *K* = 1–7 with a burn-in phase of 10,000 repetitions and a run phase of 100,000 repetitions and used default model assumptions with no weight given to location as a prior. Fisher’s exact tests were used to compare differences in estimated admixture across groups. A principal components analysis (PCA) complemented the analysis of genetic structure using the ade4 package v.1.7.13 in R (Dray & Dufour 2007).

Finally, we measured linkage disequilibrium to quantify the strength of coupling at barrier loci (*pgFAR*, *Pdfr*) and the degree to which selection and assortative mating are counteracting neutral gene flow and recombination at each of the six sites. After filtering for a 0.15 minor allele frequency cutoff, we calculated *r*^2^ values for every pair of biallelic SNPs in the female moth dataset using custom scripts in R. Following calculations, we estimated mean *r*^2^ value per locus and then combined results by barrier effect type (i.e., no barrier, one barrier, two barriers). We thinned this dataset to include only *pgFAR* and a single randomly selected locus per Z chromosome-linked scaffold. This resulted in a final dataset including 27 loci: 11 Z chromosome-linked markers outside the rearrangement (including *Pdfr*), 15 Z chromosome-linked markers within the rearrangement, and *pgFAR*. Mann-Whitney U tests were used to compare differences in LD across groups.

## RESULTS

### Pooled genome sequencing

There were no significant effects of geography on genetic differentiation, as determined by Mantel tests (for any category: “rearranged” Z, *p* = 0.47; “collinear” Z, *p* = 0.57; autosomal, *p* > 0.14; 1000 permutations), consistent with previous studies (Bourguet et al. 2000; Kim et al. 2011; Coates et al. 2019). We demonstrated qualitatively similar results with our conservative (within ~100km) analysis and could not detect any bias (Figure S2). Thus, we included all pairwise comparisons regardless of site of collection to increase the number of comparisons that could be analyzed.

All predicted chromosomes not genetically linked to trait loci showed little genetic differentiation regardless of the number of phenotypic differences between population samples (mean *F_ST_* ≤ 0.05, *p* > 0.1, Figure 2). In contrast, differentiation on the Z chromosome and chromosome 12 increased as the number of barriers accumulated. Compared to population pairs lacking phenotypic divergence (mean *F_ST_* = 0.04), differences in pheromone led to significant elevation (25%) of differentiation on chromosome 12 (mean *F_ST_* = 0.05, *p* = 0.03), whereas differences in phenology (voltinism) did not (mean *F_ST_* = 0.04, *p* = 0.86), as expected since phenology barrier loci do not occur on chromosome 12. The largest increase (50%) of chromosome 12 differentiation occurred upon joint divergence of pheromone and phenology (mean *F_ST_* = 0.06, *p* = 6.7 x 10^-5^), although the increase was not significantly greater than pheromone divergence alone (*p* = 0.17).

**Figure 2.**
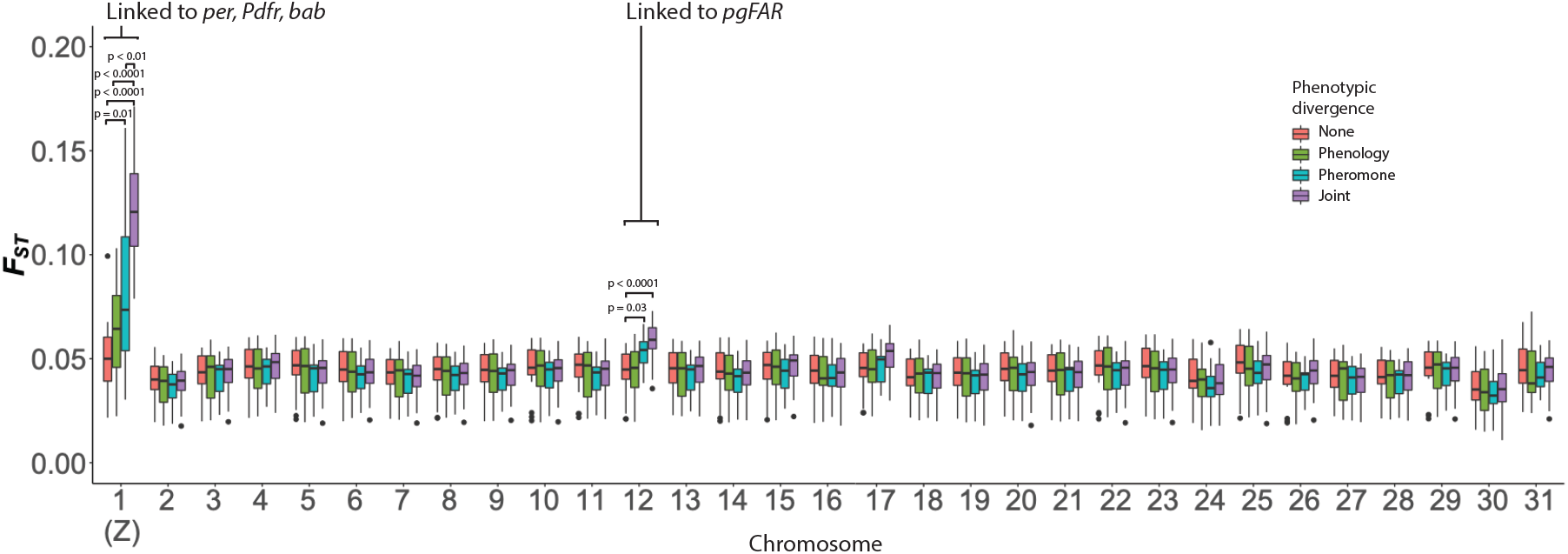
Genetic differentiation (mean *F*_ST_) estimated in 100 kb windows plotted by predicted chromosome between population pairs lacking differences in barrier traits (Z/Z; red) or differing in phenology (BZ/UZ; green), pheromone signaling (BE/BZ; blue), or both phenology and pheromone (BE/UZ; purple). Scaffolds linked to barrier trait loci *pgFAR*, *per*, *Pdfr*, and *bab* were excluded from *F*_ST_ calculations on the Z chromosome and chromosome 12. Statistically significant differences assessed via Kruskal-Wallis tests with Dunn’s test for false discovery rates are labeled with brackets.

Compared to population pairs lacking phenotypic divergence on the Z chromosome (mean *F*_ST_ = 0.05, Figure 2), phenology divergence led to a weak (20%) non-significant increase of differentiation (mean *F*_ST_ = 0.06, *p* = 0.28), whereas a significant (80%) increase was seen in the presence of pheromone divergence (mean *F*_ST_ = 0.09, *p* = 0.01). Joint divergence led to the greatest (140%) increase of Z chromosome differentiation (mean *F*_ST_ = 0.12, *p* = 5.3 x 10^-9^) and was statistically significant compared to phenology alone (*p* = 2.7 x 10^-6^) and pheromone alone (*p* = 8.2 x 10^-3^). This effect was primarily driven by elevated *F*_ST_ inside the region of reduced recombination among populations that differed in both pheromone and phenology (Figure 3D, 3E). *F*_ST_ of the rearranged region was elevated in joint divergence comparisons relative to the remaining three population categories (*p* < 0.007, Figure 3D). *F*_ST_ of collinear regions was significantly elevated between populations with joint divergence compared to populations lacking trait differences and populations differing only in phenology (*p* < 0.007, Figure 3E). However, *F*_ST_ was not quite significantly elevated compared to pairs that differed only in pheromone (*p* = 0.11). In populations that differed in just pheromone, *F*_ST_ inside the region of reduced recombination was elevated compared to populations lacking barrier effects (*p* = 0.007), but not populations differing in phenology (*p* = 0.23, Figure 3D).

**Figure 3.**
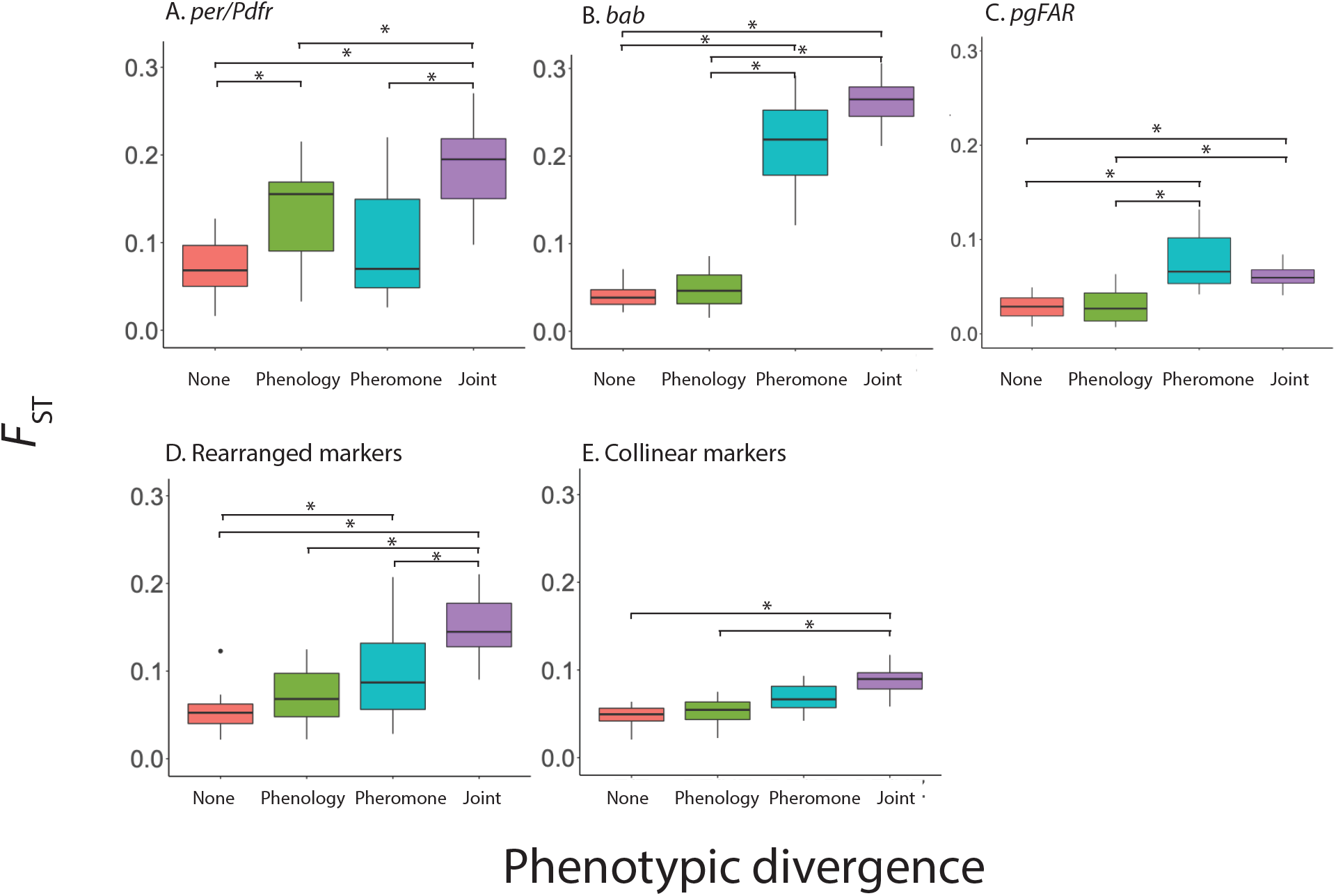
A. Genetic differentiation (*F*_ST_) at phenology barrier loci *per* and *Pdfr* between population pairs lacking differences in barrier traits (Z/Z; red) or differing in phenology (BZ/UZ; green), pheromone signaling (BE/BZ; blue), or both phenology and pheromone (BE/UZ; purple). B. Genetic differentiation at pheromone response barrier locus *bab* across population pairs. C. Genetic differentiation at pheromone production barrier locus *pgFAR* across population pairs. D. Genetic differentiation at neutral Z chromosome loci inside a ~10 Mb “rearranged” region of suppressed recombination across population pairs. E. Genetic differentiation at neutral Z chromosome loci outside a ~10 Mb region of recombination suppression (“collinear” region) across population pairs. Asterisks indicate significantly different comparisons following Kruskal-Wallis tests and Dunn’s tests accounting for false discovery rates (corrected p < 0.05).

With respect to barrier loci, the phenology loci (*per/Pdfr*) had significantly elevated *F*_ST_ in jointly diverged population pairs compared to population pairs without phenology differences (*p* < 0.0016), as well as those differing only in phenology (*p* = 0.017, Figure 3A). *F*_ST_ in the *bab* region significantly increased in all populations that differed in pheromone compared to populations that did not (*p* < 0.0003, Figure 3B), but *F*_ST_ was not quite significantly increased upon joint divergence compared to pheromone alone (*p* = 0.11). *F*_ST_ in the *pgFAR* region was broadly similar to that seen at *bab* across population pairs. The 100 kb window with the highest mean *F_ST_* on the Z chromosome in populations differing only in phenology overlapped the voltinism gene *per* (scaffold532: *F*_ST_ = 0.23; Figure S4), in populations differing only in pheromone signaling it was within the *bab* region (scaffold50: *F*_ST_ = 0.39), and the highest window in jointly diverged populations was inside the region of recombination suppression (scaffold804: *F*_ST_ = 0.61).

### Targeted amplicon sequencing

The number of distinct genotypic clusters and degree of admixed ancestry varied across localities by the number of acting barriers. The most likely number of clusters across populations was two for all STRUCTURE runs (*K* = 2, Evanno et al. 2005). The two locations lacking any known barrier effect (Table 1) showed strong assignment to a single cluster. For all 73 individuals, more than 98% of their Z chromosome loci belonged to the same cluster (Qs > 0.98; Figure 4A). The two locations differing in pheromone but not phenology (BE/BZ) showed evidence of two genotypic clusters and greater admixture. At these sites, 22% (19 of 83) of individuals showed admixture (defined as 0.1 < Q < 0.9). The two locations where pheromone and phenology could act coincidentally (BE/UZ) also showed two genotypic clusters, but unlike at sites where insects differed only in pheromone, most individuals showed limited admixture of the Z chromosome and the proportion of admixed individuals was reduced. 96% of individuals at BE/UZ sites had a high probability of belonging to either cluster 1 (N = 21, Q1 ≥ 0.9) or cluster 2 (N = 37, Q2 ≥ 0.9).

**Figure 4.**
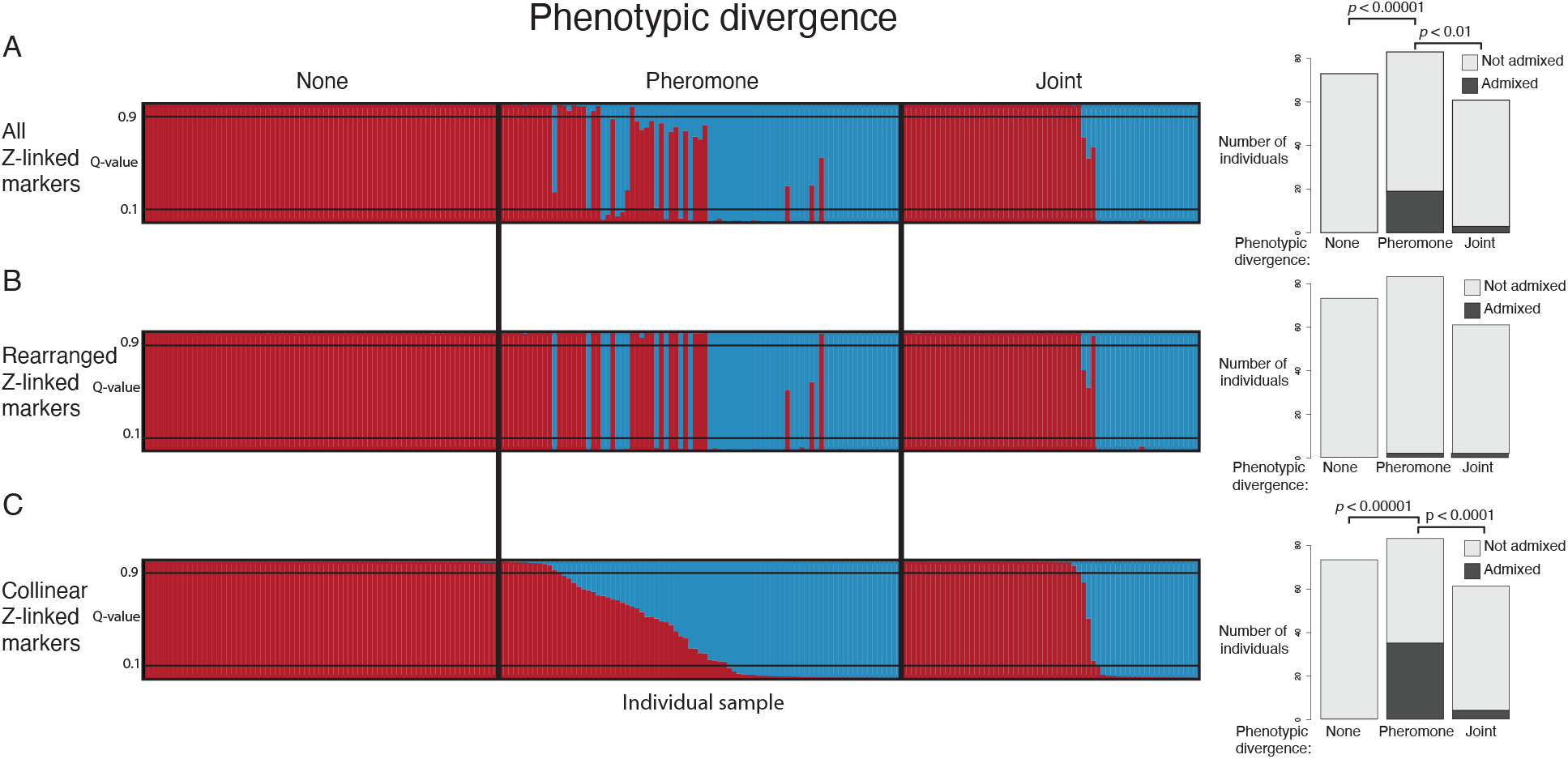
STRUCTURE plots partitioned by population category. Plotted on the right are the number of individuals that are considered admixed (Q-value greater than 0.1 and less than 0.9) and not-admixed (Q value greater than 0.9 or less than 0.1). A. All Z chromosome markers; B. rearranged Z markers; C. collinear Z markers. Red indicates assignment to cluster 1 (Z clade), blue indicates assignment to cluster 2 (E clade). Comparisons are between populations lacking differences in barrier traits (None; Z/Z), or pheromone signaling (Pheromone; BE/BZ), or both phenology and pheromone (Joint; BE/UZ).

Separate analyses of Z chromosome loci inside and outside the rearranged region showed that admixture depended on marker location (Figure 4), in addition to barriers to gene flow. Population types significantly differed in the degree of admixed collinear regions (*p* = 7.63 x 10^-14^), with an excess of admixture at sites with insects differing only in pheromone compared to sites without phenotypic divergence or sites with joint differences in phenology and pheromone. Admixture of rearranged regions was similarly low across all three population types (*p* = 0.386). At sites where insects differed only in pheromone, nearly half (42%) of 83 individuals were estimated to have admixed collinear loci (0.1 < Q < 0.9), whereas only 2 showed appreciable admixture at loci in the rearranged region (*p* = 1.64 x 10^-10^). In contrast, at sites differing in both pheromone signaling and phenology, both collinear and rearranged regions were relatively free of admixture and the estimated proportion of admixed Z chromosomes did not significantly differ (3.3% vs. 6.6%, *p* = 0.68). Consistent with a lower per generation rate of admixture across the rearrangement, a large percentage of individuals of mixed ancestry inferred from the rearranged region resembled F_1_ hybrids (Q = 0.4 – 0.6; 3 of 4 admixed, 75%) compared to regions with normal recombination rates (11 of 39 admixed, 28.2%) (*p* = 0.094). The former may represent F_1_ heterokaryotype male offspring.

We evaluated Z chromosome admixture relative to allelic associations at the autosomal gene *pgFAR* and found that sex-linked genotypic clusters 1 and 2 mainly corresponded to E and Z pheromone alleles, respectively. Moths assigned with high probability to either of the two clusters (Q1 or Q2 ≥ 0.9, N = 168) were usually homogenous for E or Z alleles at *pgFAR* (145; 86%), whereas 25 percent of moths showing evidence of admixture (0.1 < Q < 0.9) had heterozygote *pgFAR* SNP genotypes. We used a simple network to visualize the E genotypic cluster (Q1 > 0.9), the Z genotypic cluster (Q2 > 0.9), and the admixed cluster (0.1 < Q1 < 0.9) separated by *pgFAR* genotype (Figure 5). Most moths demonstrating a mismatch between their autosomal *pgFAR* genotype and their genotypic cluster assignment (N = 48) were collected at sites differing only in pheromone (40 of 77) (Figure 5). In contrast, sites differing in both pheromone and phenology had fewer discordant individuals (8 of 43). Results from the PCA of the Z chromosome (Figure 6, Figure S3) were consistent with these results. The first coordinate axis was broadly consistent with *pgFAR* assignment (Figure S3), and whereas intermediates in coordinate space of an analysis of collinear markers were primarily admixed individuals at BE/BZ sites (Figure 6A), few intermediates occurred when analyzing solely rearranged loci (Figure 6B).

**Figure 5.**
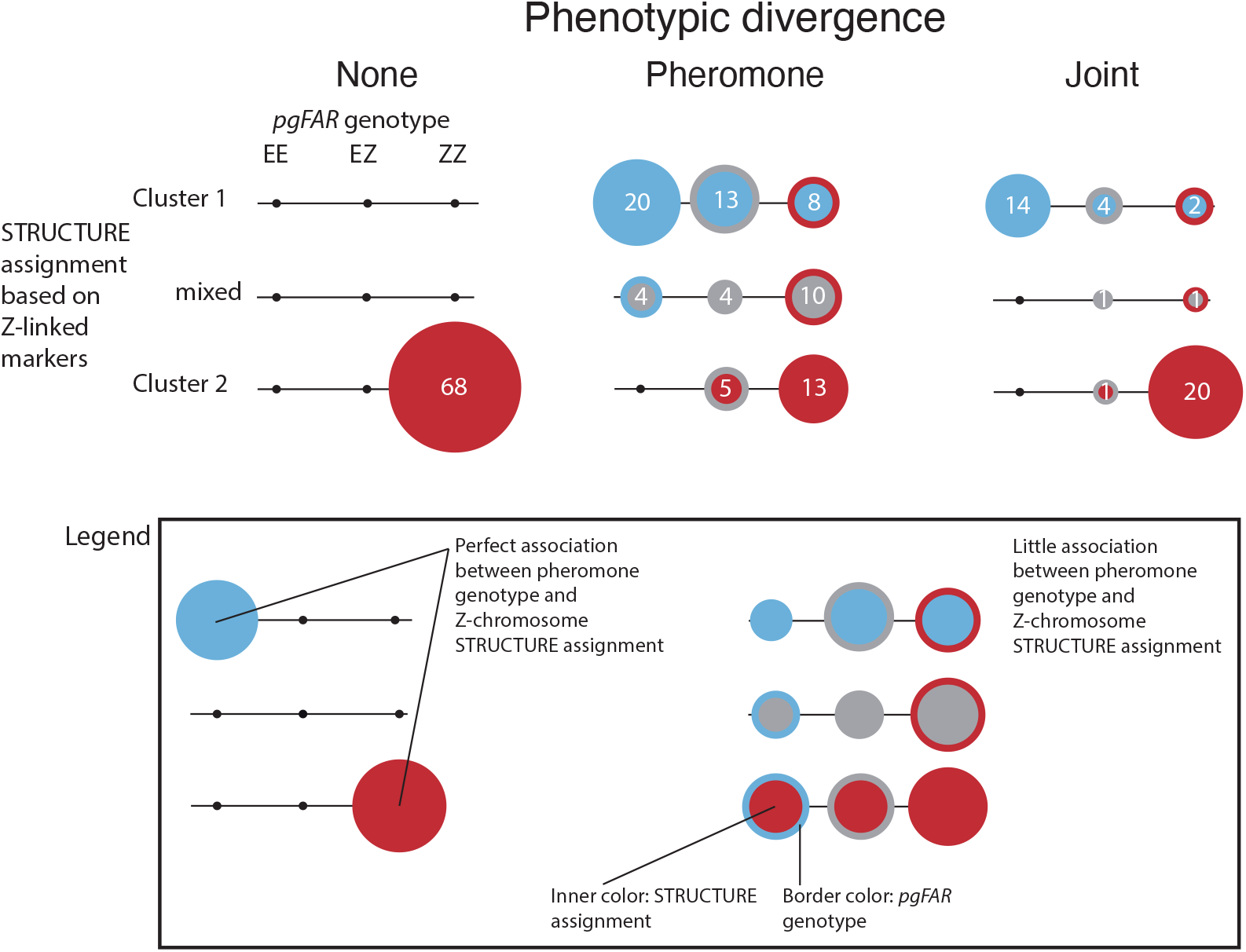
The accumulation of barrier traits leads to increase concordance between *pgFAR* and STRUCTURE clade assignment. Bead-and-string plots depict association between sample genotypes at *pgFAR*, the autosomal locus controlling pheromone production (border color) and Z chromosome marker-based STRUCTURE assignment (inner color). Samples were considered E-clade if they were assigned to an E-strain associated STRUCTURE background at a 0.9 threshold or above, mixed if between 0.1 and 0.9, and Z-clade if below 0.1 (see Figure 4). Number of individuals listed inside beads. Comparisons are between populations lacking differences in barrier traits (None; Z/Z), or pheromone signaling (Pheromone; BE/BZ), or both phenology and pheromone (Joint; BE/UZ).

**Figure 6.**
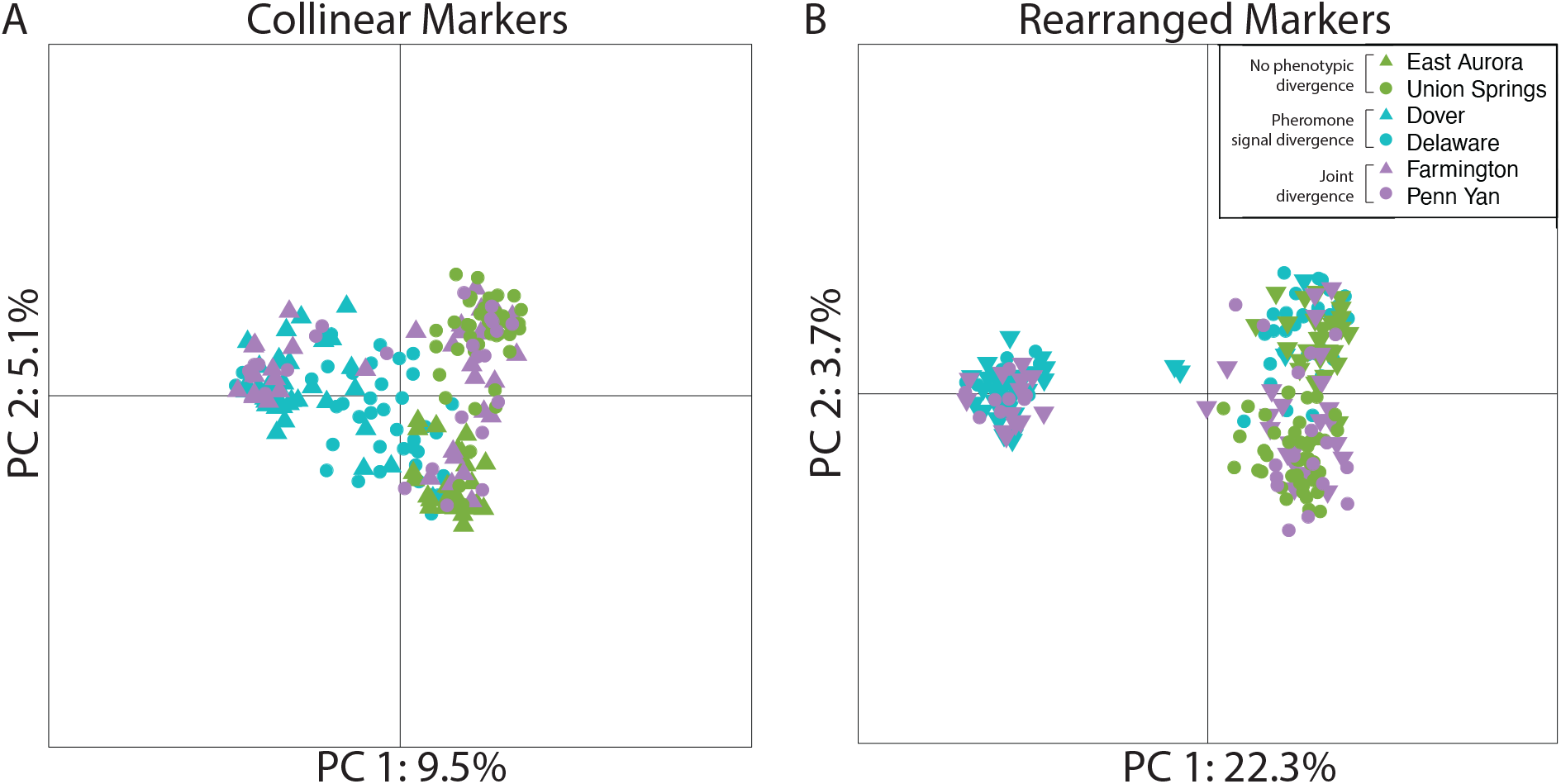
Principal components analyses of (A) collinear and (B) rearranged Z chromosome markers. Individual labels are based on geographic location and population type. Percentages indicate the amount of variance explained by that axis. Comparisons are between populations lacking differences in barrier traits (Z/Z; green), or pheromone signaling (BE/BZ; blue), or both phenology and pheromone (BE/UZ; purple).

LD across the entire Z chromosome increased as the number of barrier traits increased. *r*^2^ values were low at locations without barriers (mean *r*^2^ =0.027), but progressively grew with the presence of a single barrier (~120%) (BE/BZ, mean *r*^2^ = 0.06, *p* = 1.00 x 10^-20^) and coincident barriers (~310%) (BE/UZ, mean 0.112, *p* = 5.64 x 10^-54^) (Figure 7). Finally, LD between *pgFAR* and *Pdfr* increased primarily upon joint divergence, as did LD between *pgFAR* and the rest of the Z chromosome (Figure 8). Splitting loci by collinear and rearranged location showed that the influence of barrier effects on LD were more pronounced in rearranged intervals (Figure S5).

**Figure 7.**
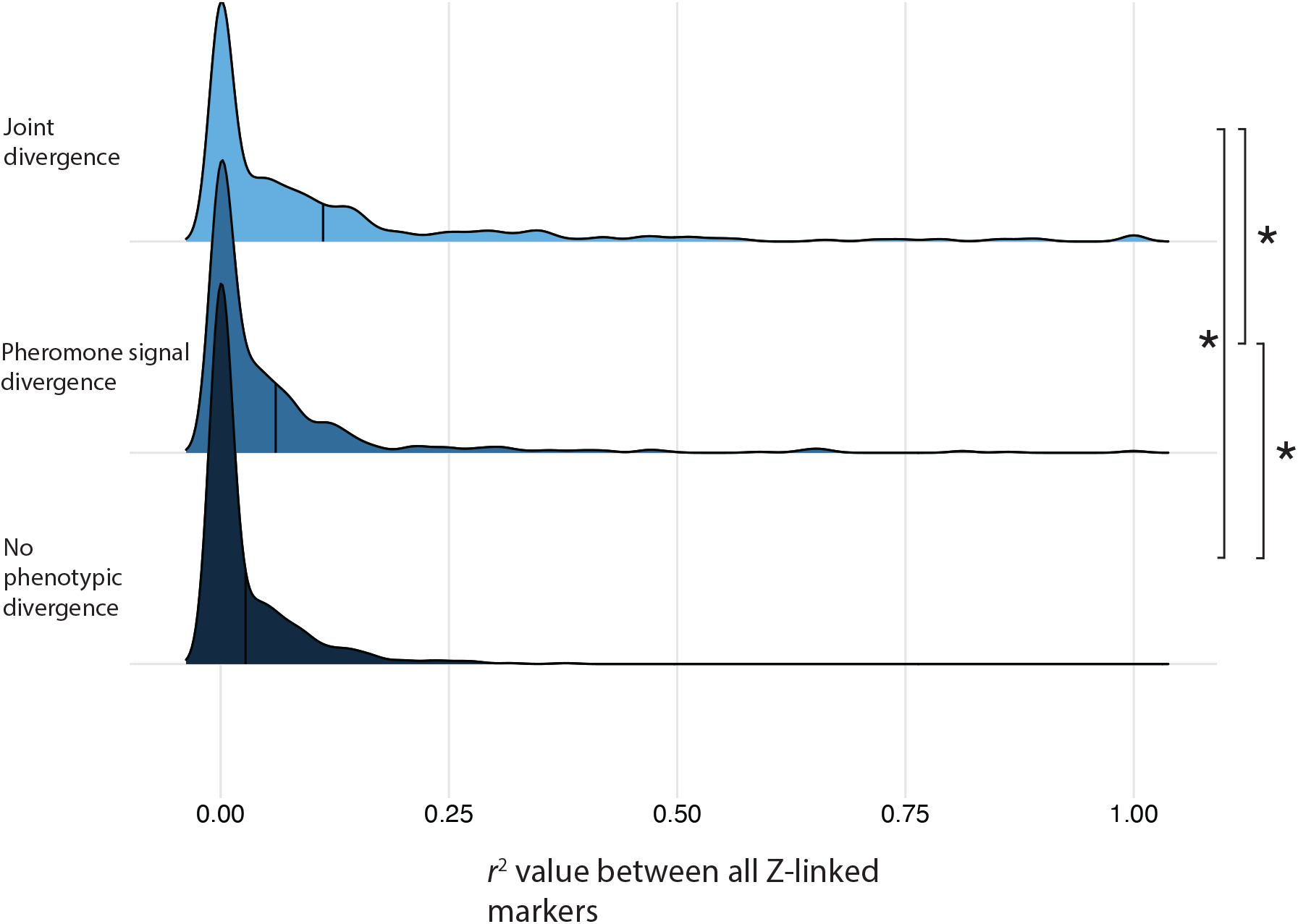
Linkage disequilibrium (*r*^2^) between all Z chromosome SNPs separated by population type. The mean value is indicated by a vertical black line. Significantly different distributions measured using Mann-Whitney U tests are marked with asterisks (*). Comparisons are between populations lacking differences in barrier traits (Z/Z), or pheromone signaling (BE/BZ), or both phenology and pheromone (BE/UZ).

**Figure 8.**
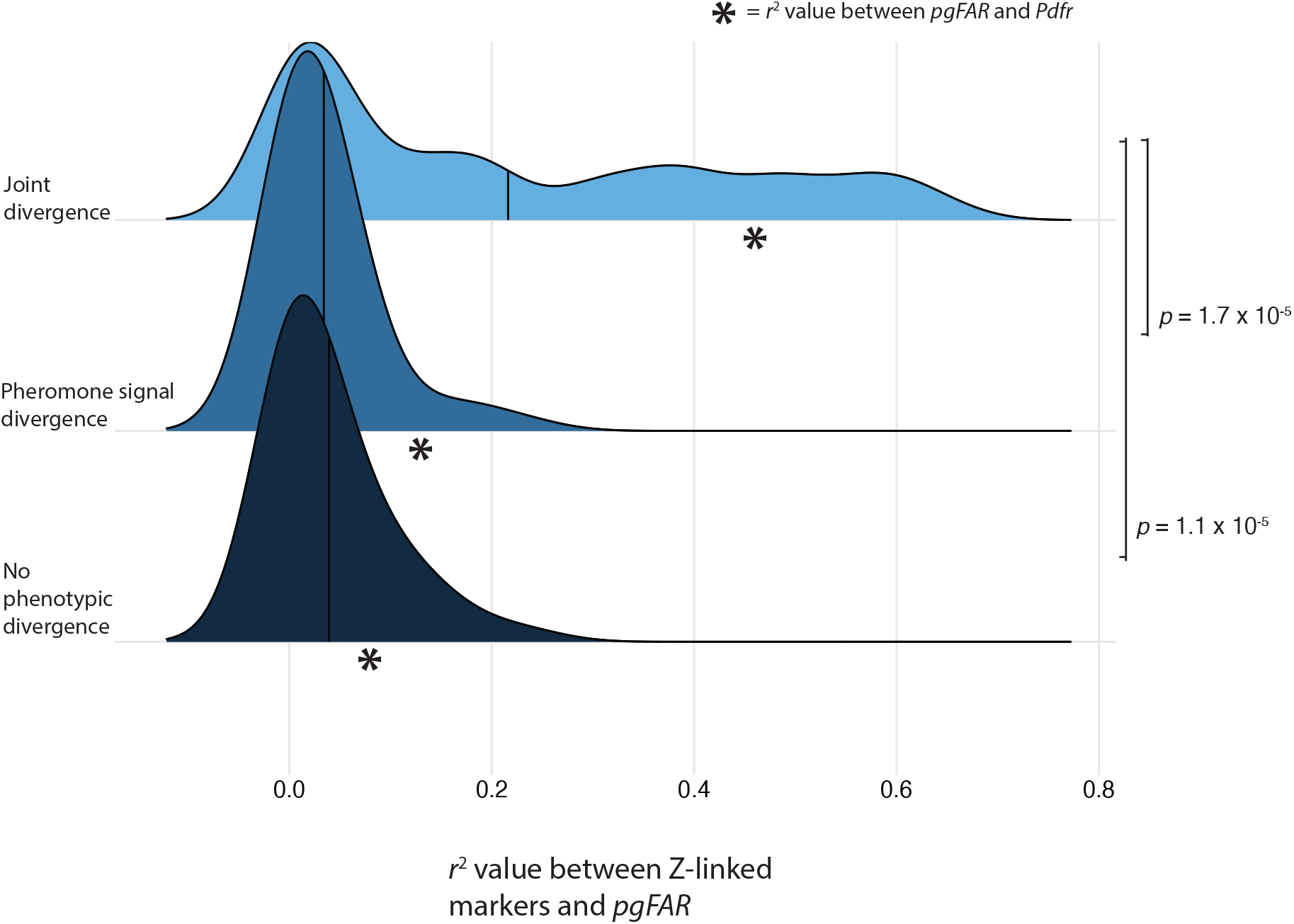
Linkage disequilibrium (*r*^2^) calculated between the autosomal *pgFAR* gene and SNPs on the Z-chromosome, separated by population type. The mean value is indicated by a vertical black line. The star depicts mean LD between SNPs at *pgFAR* and the Z-linked *Pdfr* gene. Significance from Mann-Whitney U tests comparing population types are shown. Comparisons are between populations lacking differences in barrier traits (Z/Z), or pheromone signaling (BE/BZ), or both phenology and pheromone (BE/UZ).

**Figure 9.**
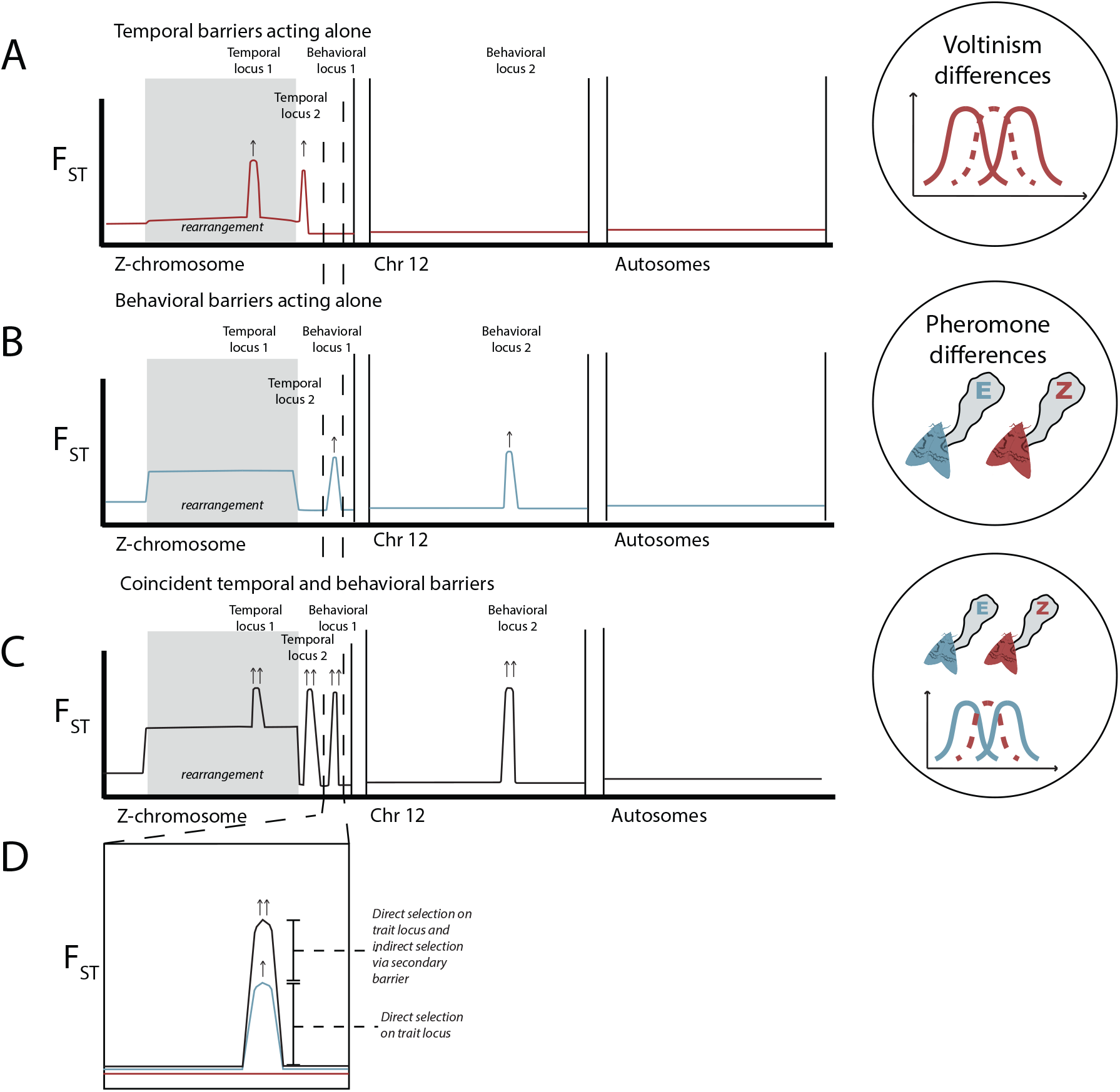
A graphical overview of genomic differentiation as barriers to gene flow accumulate. Patterns reflect key findings but are not real data. A. Differentiation at locations differing only in phenology (temporal barrier), B. only in pheromone signaling (behavioral barrier), or C. joint divergence along both phenotypic axes (temporal and behavioral barriers). D. An illustration of how coincidence shapes differentiation of barrier loci by allowing both direct and indirect selection.

## DISCUSSION

The origin of distinct genotypic clusters as speciation progresses is thought to depend on 1) the number and genomic architecture of segregating barrier loci, 2) strength of selection or assortative mating on each locus, and 3) features that reduce the rate of recombination among barrier loci (Butlin et al. 2021, Felsenstein 1981, Nosil et al. 2017, Seehausen et al. 2014). The ECB moth system is interesting because many of these factors are at least partially understood. Genomic locations of barrier loci and a Z-chromosomal rearrangement are known, the strength of relevant barriers have been estimated, and barrier effects among populations can be independent or coincident, allowing insight into how the transition from a weaker barrier(s) to a stronger overall barrier relates to the emergence of exclusive genetic groups.

### Phenology divergence alone

Our findings indicate that a >20 day shift in phenology between univoltine Z (UZ) and bivoltine Z (BZ) populations results largely in regions of differentiation overlapping Z-linked *per* and *Pdfr* loci controlling the timing of diapause termination (Figure 2, 3A, 3C, 3D). Lack of widespread genetic differentiation was somewhat unexpected because the estimated strength of assortative mating using Coyne and Orr’s (1989) method indicates intermediate reproductive isolation (mean *RI* = 0.66, Dopman et al. 2010). Moreover, as a ‘magic trait,’ reproductive isolation would be enhanced if hybrid individuals suffer low fitness because their intermediate phenology causes a temporal mismatch with the seasonal environment (Kozak et al. 2017; 2019; Servedio et al. 2011). Among interacting populations, the chromosomal rearrangement spanning ~10 Mb of the Z-chromosome (from ~6 Mb to ~16 Mb) (Wadsworth et al. 2015; Kozak et al. 2017) might conceivably help reduce the number of such intermediate genotypes, as it includes *per* located at ~13.5 Mb (but not *Pdfr* at ~18 Mb) (Kozak et al. 2019; Unbehend et al. 2021). Yet we found limited evidence of enhanced differentiation in this part of the Z chromosome (Figure 3C), which would have been expected if a rearrangement differed in frequency and was helping maintain high fitness genotypes by preventing recombination (Kirkpatrick & Barton 2006).

Shallow phenology-associated genetic structure is also seen in other insect systems, for example summer and winter forms of the pine processionary moth (Santos et al. 2007, 2011), as well as marine midge populations living in different phenological niches (Fuhrmann et al. 2021). One possible explanation for weak, localized genetic isolation in these systems is if temporal barriers are commonly conditionally dependent, and therefore unstable. Like other insect species, year-to-year differences in weather alters breeding cycle phenology in the ECB moth (Beck 1983), which can lead to occasional increased overlap of allochronic mating flights or possibly ease fitness consequences of intermediate hybrid phenology, thereby allowing genomic regions not tightly linked to selected barrier loci to be exchanged. In possible agreement with conditional dependence, Dopman et al. (2010) found the barrier effect to vary by as much as two-fold across years. Despite the potential for divergent selection on phenology to efficiently reduce gene flow, allochrony does not therefore appear to be strong enough to promote divergence by itself, a question recently considered across a variety of taxa (Taylor and Friesen 2017). Instead it may more commonly function when combined with other barriers, such as habitat or resource partitions (Doellman et al. 2018), or when reinforcing pre-existing divergence (Lowry et al. 2008; Møller et al. 2011).

### Behavioral divergence alone

There is general agreement that many of the ~160,000 moth species remain separate because of changes in pheromone preference and pheromone blend. However, as with ongoing discussions on phenology, current debate centers on whether this phenotypic axis suffices to initiate the origin of new species (reviewed in Allison and Cardé 2016). The buildup of genome-wide differentiation that ultimately characterizes species is thought to be favored by ‘multiple-effect traits’ like pheromone signaling in ECB moths, because they are highly efficient at reducing gene flow (Smadja and Butlin 2011). Indeed, the two routes of reproductive isolation that evolved from pheromone divergence (mate choice and behavioral sterility) are predicted to eliminate almost 90% of gene flow between Z and E strains, at least under laboratory conditions (mean *RI* = 0.89; Glover et al. 1991; Dopman et al. 2010). In the field, Unbehend et al. (2021) recently showed that male moths carry either E type or Z type alleles at *bab* (pheromone preference) and at *pgFAR* (pheromone blend) genes ~80% of the time, indicating coupled pre- and post-zygotic isolation is strong enough in nature to counteract randomization of these physically unlinked genes.

Our findings corroborate prior inferences of selection and assortative mating on *bab* and *pgFAR* barrier loci in field populations (Figure 3B, 3C), but they go on to suggest that divergence of pheromone signaling alone may not necessarily lead to the emergence of genome-wide differentiation (Figure 2). Nevertheless, bivoltine E (BE) and bivoltine Z (BZ) populations do exhibit broad differentiation of chromosomes harboring *bab* and *pgFAR* genes (Z chromosome and autosome 12) (Figure 2, 3D, 3E), as well as long-range LD between sympatric populations (Figure 7). Differentiation was unexpectedly high within rearranged regions of the Z chromosome (Figure 3D, 3E), despite *bab* (~18.7 Mb) being collinearly located ~3 Mb away from the nearest estimated rearrangement breakpoint (Figure S1). This might indicate that divergent selection acts on another component of the pheromone system controlled by loci within the rearrangement. If true, two prime suspects are the tandemly repeated odorant receptor loci *OR4* and *OR6* (~12.14 Mb) required for physiological perception of E and Z pheromone components within the male antenna (Koutroumpa et al. 2014). Overall, the apparent stability of chromosome but not genome-wide differentiation implies the sex pheromone system of moths might reduce gene flow enough to allow a partial build-up of differentiation, but this phenotypic axis alone may not lead to substantial progress towards the emergence of well-defined species.

### Joint phenotypic divergence

Our analyses of bivoltine E (BE) and univoltine Z (UZ) moths indicate that joint divergence along multiple phenotypic axes (phenology and pheromone signaling) is associated with intensified genetic differentiation. The expected progression towards enhanced coupling as behavioral and temporal barriers coincide in sympatry was detected as elevated LD between unlinked behavioral barrier and temporal barrier genes (*pgFAR* and *Pdfr*) (Figure 8). Differentiation at temporal barrier loci (*per*, *Pdfr*) increased in the presence of behavioral divergence (Figure 3A, phenology vs. joint divergence). Likewise for the behavioral barrier locus *bab* in the presence of phenology divergence (Figure 3B, pheromone vs. joint divergence). Unlike *bab*, differentiation was not elevated at *pgFAR* (Figure 3C, pheromone vs. joint divergence), a result likely explained by exclusion of *pgFAR* heterozygotes when creating phenotypically divergent population pools for sequencing. As shown by the amplicon data (Figure 5), *pgFAR* heterozygotes are more than twice as frequent at sites differing only in pheromone signaling (BE/BZ), compared to sites that also differ in phenology (BE/UZ). Therefore their removal may have artificially inflated estimated *pgFAR* genetic differentiation in BE/BZ comparisons (e.g., Landisville, Dover). In contrast, levels of differentiation at this locus in comparisons of jointly diverged sites may be more reflective of reality, since heterozygotes are relatively rare (e.g., Rockspring, Harner).

The wider genomic landscape appears to be similarly influenced by joint divergence. On the Z chromosome, both long-range LD (Figure 7) and inter-chromosomal LD with *pgFAR* (Figure 5, 8, S4) increased upon the coincidence of behavioral and temporal barriers in sympatry. Furthermore, whereas evidence of reduced admixture (or greater clustering) on the Z was mainly limited to the rearrangement when sympatric populations differed in pheromone signaling, this signature expanded into collinear regions when sympatric populations also differed in phenology (Figure 4, Figure 6). Finally, overall differentiation of the two chromosomes containing the four barrier loci (Z chromosome and autosome 12) increased by more than two-fold (Figure 2), although differentiation is disproportionate on the Z chromosome where coupling leads to a ‘large Z-effect’ and divergence of three barrier loci (*per–Pdfr–bab* complex), compared to just one on chromosome 12 (*pgFAR*).

All these features are congruent with prior estimates of cumulative reproductive isolation. When differences in sex pheromone communication as well as phenology act as barriers, their cumulative effects are estimated to result in almost complete reproductive isolation (mean *RI* = 0.96, Dopman et al. 2010). The process underlying heightened genetic differentiation is unlikely to simply involve a sudden increase in fitness effect on each barrier locus (direct selection). Instead, as barrier loci increase their non-random associations during coupling, a higher mean fitness differential can be achieved between alternative multi-locus genotypes (indirect selection). Since each locus can have its own fitness effect and also be indirectly influenced by the fitness effect of other loci (Barton 1983), overall selection is more effective at each locus, as is the overall barrier to gene flow at each locus, making it more difficult for neutral alleles to cross population boundaries (Barton and Bengtsson 1986).

### Conclusions

We argue that our study provides support for the coupling hypothesis of speciation (Felsenstein 1981; Smadja and Butlin 2011; Seehausen et al. 2014) and helps close a gap between theoretical predictions of this hypothesis and empirical systems. Our population comparisons show that whether barriers are independent or coupled make important contributions to a hallmark of the speciation process, the build-up of genomic differentiation. Genetic distances fit expectations from estimates of reproductive isolation, with locus-specific differentiation associated with the weakest barrier (temporal), differentiation across much of a chromosome with the barrier of moderate strength (behavioral), and chromosome-wide differentiation when the two barriers operate cumulatively. These results demonstrate that the joint action of multiple, coincident barrier effects lead to levels of genomic differentiation that far exceed those of single barriers acting alone, consistent with theory in which coupling allows indirect selection to combine with direct selection and thereby result in a stronger overall barrier.

Finding chromosome-wide but not genome-wide differentiation when populations diverge along multiple phenotypic axes suggests that the overall barrier upon coupling may only partly outweigh gene flow and recombination. Thus, a critical tipping point or threshold of coupling that could allow for sudden differentiation of most of the genome was not reached (Nosil et al. 2017). When barriers are coincident, much of the genome appears well mixed and moderate genetic differentiation (*F*_ST_ = 0.05 – 0.15) is observed for only two chromosomes containing barrier loci, making further divergence challenging without evolution of either more barrier loci or stronger selection/assortative mating. Simulations demonstrate that, in certain cases, many divergently selected loci may be required for genome-wide differentiation (Feder et al. 2014, Flaxman et al. 2014). Some empirical studies seem to support these predictions, including Midas cichlids, where a transition from locus-specific to genome-wide divergence coincides with a transition from simple to polygenic barriers (Kautt et al. 2020). Strengthening of the overall barrier in the ECB moth could in time be facilitated by the evolution of highly polygenic traits, perhaps in the form of intrinsic post-zygotic barriers. However, it is also possible that many intermediate stages of speciation seen in nature, including the ECB moth, are either stably maintained or even temporary, and that barrier traits may not remain coincident should ecological or geographic shifts take place (Seehausen 2006). If so, then coupling and current population differentiation will disappear or rebound according to external environmental factors.

Understanding how and why barrier traits evolve and become coincident or disappear remains as a critical problem for speciation biology, and few empirical systems have been able to directly assess coincidence of known barrier traits in different stages of coupling. Future work on this study system will reconstruct the evolutionary and demographic history of this group and provide a framework for inferring the timing and selective history of genes underlying barrier phenotypes. Ultimately, we must reconstruct not only the sequence of evolution of individual barriers, but also the timing and causal factors responsible for their coupling. Although byproduct explanations have been proposed that do not require coupling itself to be selectively favored (Butlin and Smadja 2017), adaptive coupling seems likely in the ECB moth and can be experimentally studied by measuring fitness of full allelic combinations in nature. Indirect approaches are equally promising for identifying factors associated with changes of coupling in nature. Since it is now possible in this system to precisely estimate population differentiation and the extent of coupling of all major barrier loci (*bab*, *pgFAR*, *per*, *Pdfr*) across various spatial, ecological, and historical contexts, the causal links that establish barrier coupling and facilitate neutral divergence during speciation appear to be within reach.

## Supporting information

All_supplemental_figures_and_tables

## Notes

The authors declare that there are no conflicts of interest pertaining to this study.

### Competing Interest Statement

The authors have declared no competing interest.

### Summary of Updates

Text was edited throughout to reduce total word count. One paragraph in the conclusions section was removed, as it was part of an extended discussion and we wanted to reduce total word count for submission to a journal. Figure 7 was moved to the supplementary files and replaced with a similar figure that focuses on the entire chromosomal pattern, rather than subdivisions between collinear and rearranged regions. Typo corrected on line 164. January 2022 changes below Updated the cover page and author list to correct and authors name and added a conflict of interest statement. Corrected various typos throughout the paper and added citations throughout. Adjusted figure 1 for an error. Added version numbers to software. Moved some text from results to methods. Corrected minor error in figure 2. Updated some p-values following revisions. No interpretations changed. Added a missing significance bar in figure 3. Changed clarity of figure 5.

