## Supplementary material for "Consequences of coupled barriers to gene flow for the build-up of genomic differentiation": All_supplemental_figures_and_tables

### SUPPLEMENTARY FIGURES AND TABLES

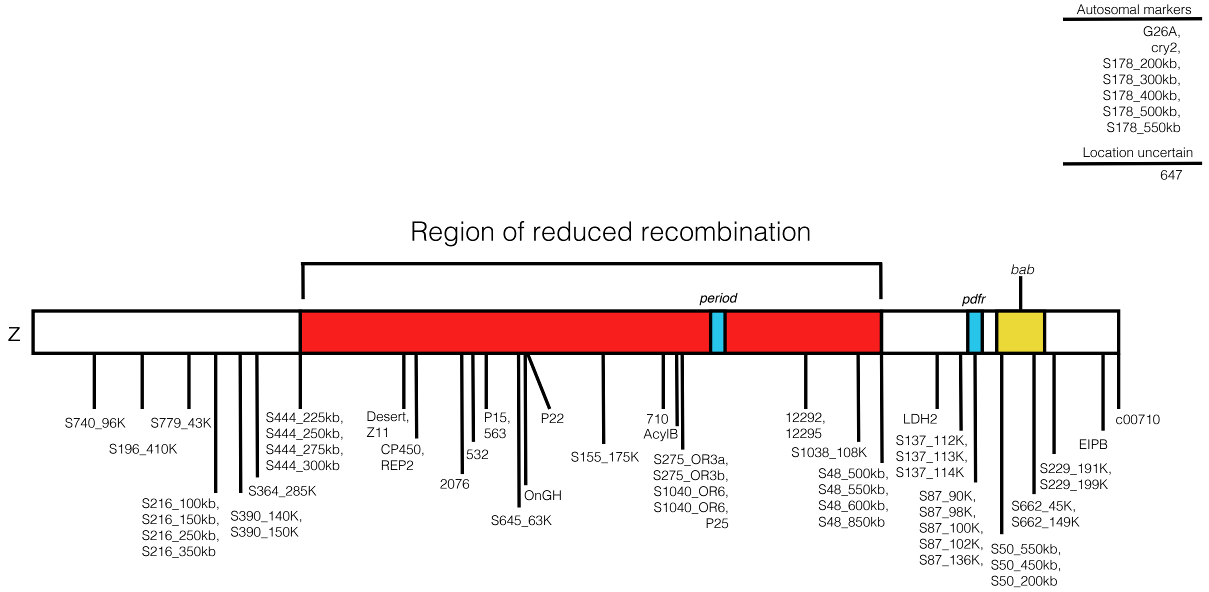


Figure S1. Map of marker locations on the Z-chromosome. Indicated are the large region of reduced recombination, *per* and *Pdfr*, loci responsible for seasonal timing and the male pheromone response locus *bab*.

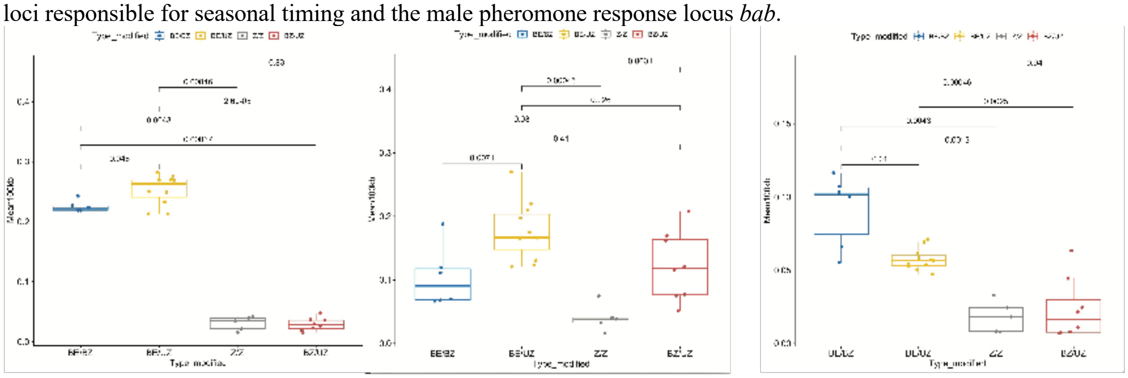

Figure S2. Pairwise $F_{ST}$ comparisons of barrier loci restricted to populations within ~100km, or roughly 1-3 generations of dispersal.

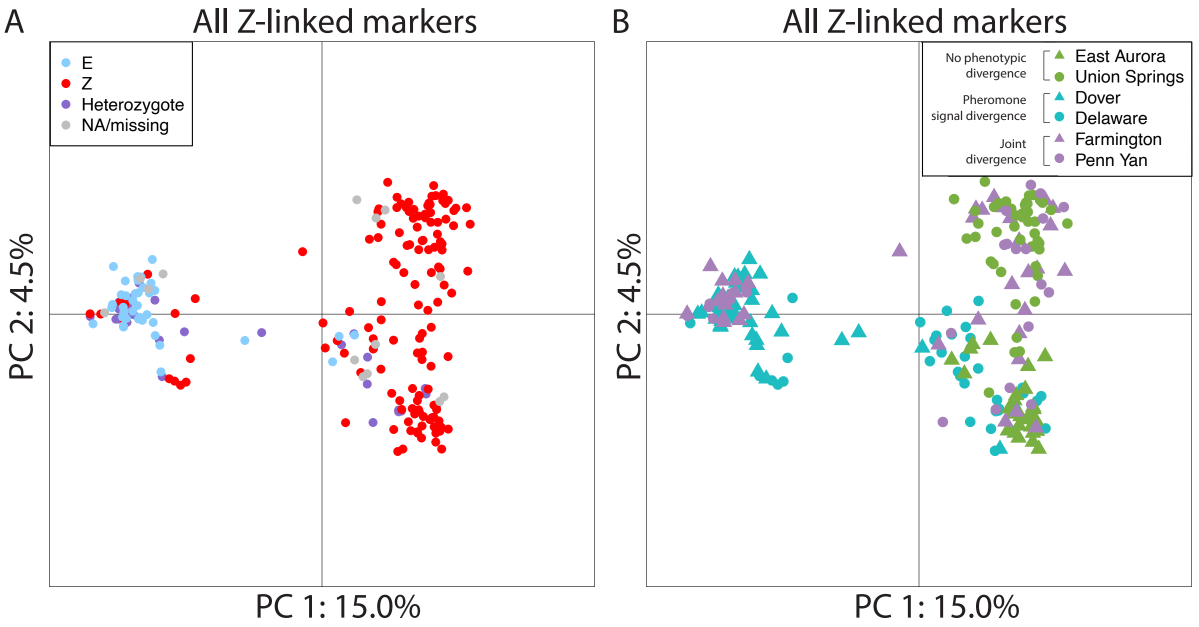


Figure S3. Principal components analyses based on Z-linked marker datasets. On the left, individual labels are based on *pgFAR* genotype. On the right, individual labels are based on geographic location and population type. Percentages indicate the amount of variance explained by that axis.

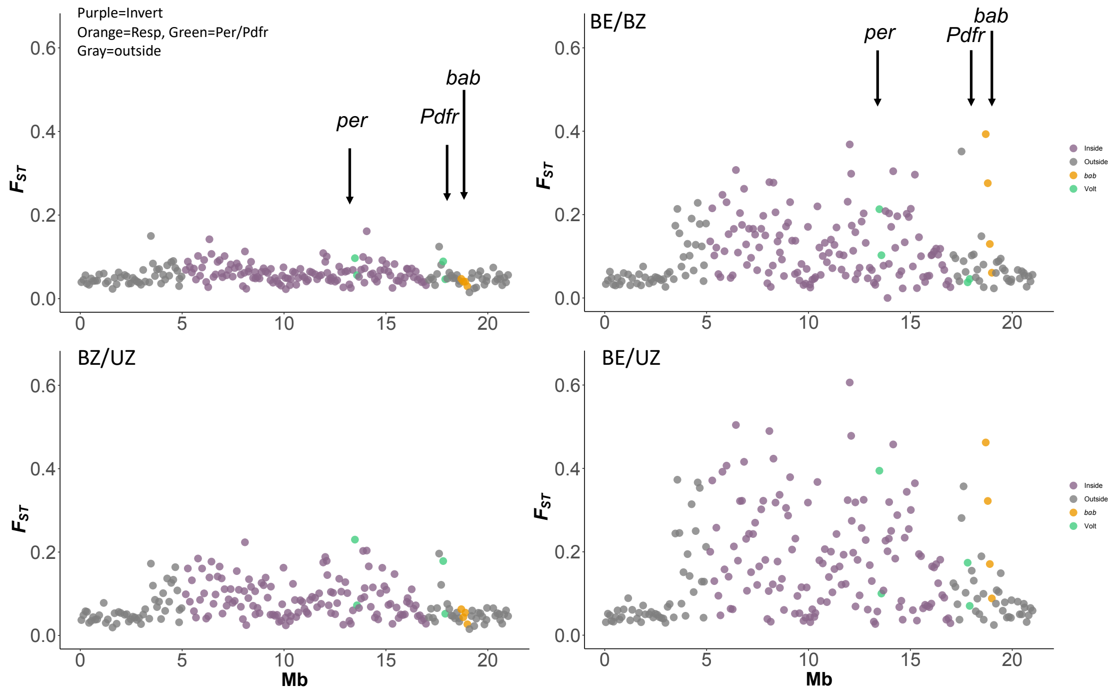


Figure S4. *F*st 100kb windows across the Z-chromosome using pooled sequencing data for four population type comparisons. Colors distinguish the rearrangement (purple) and key barrier traits (teal and orange).

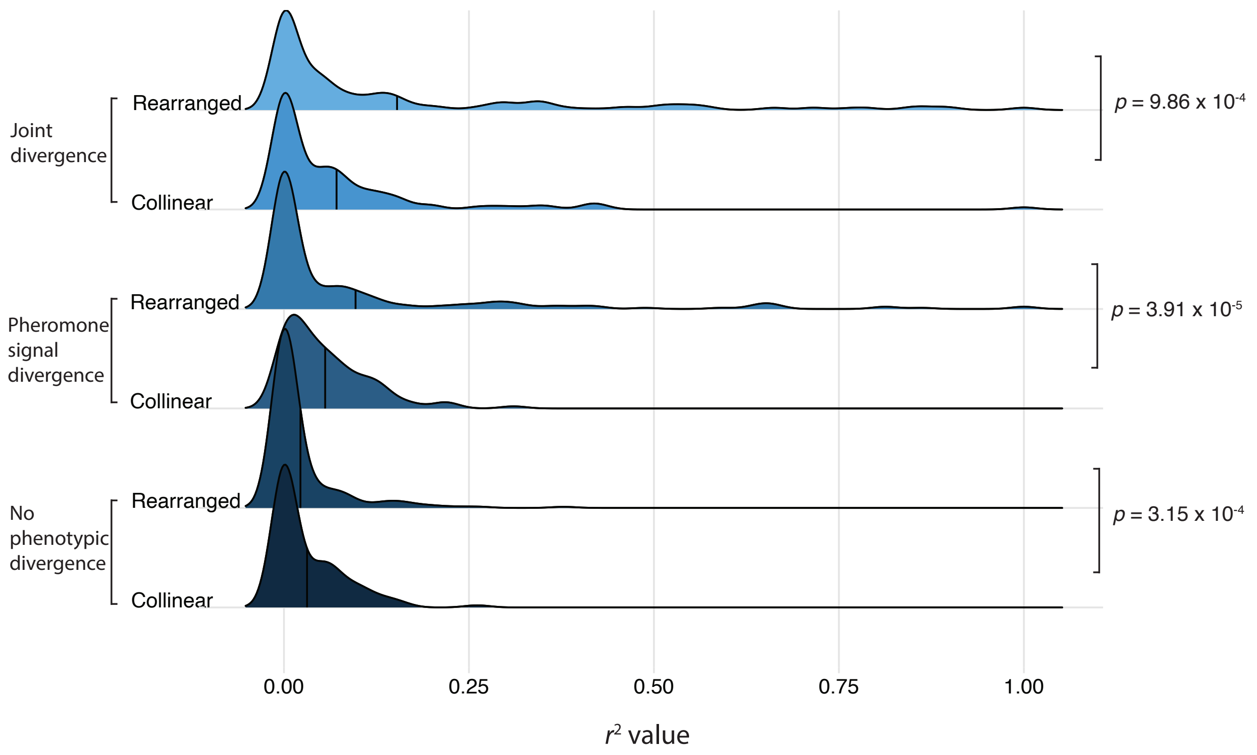


Figure S5. Linkage disequilibrium ($r^2$) between all Z chromosome SNPs separated by population type. The mean value is indicated by a vertical black line. Significance from Mann-Whitney U tests comparing rearranged and collinear $r^2$ is shown.

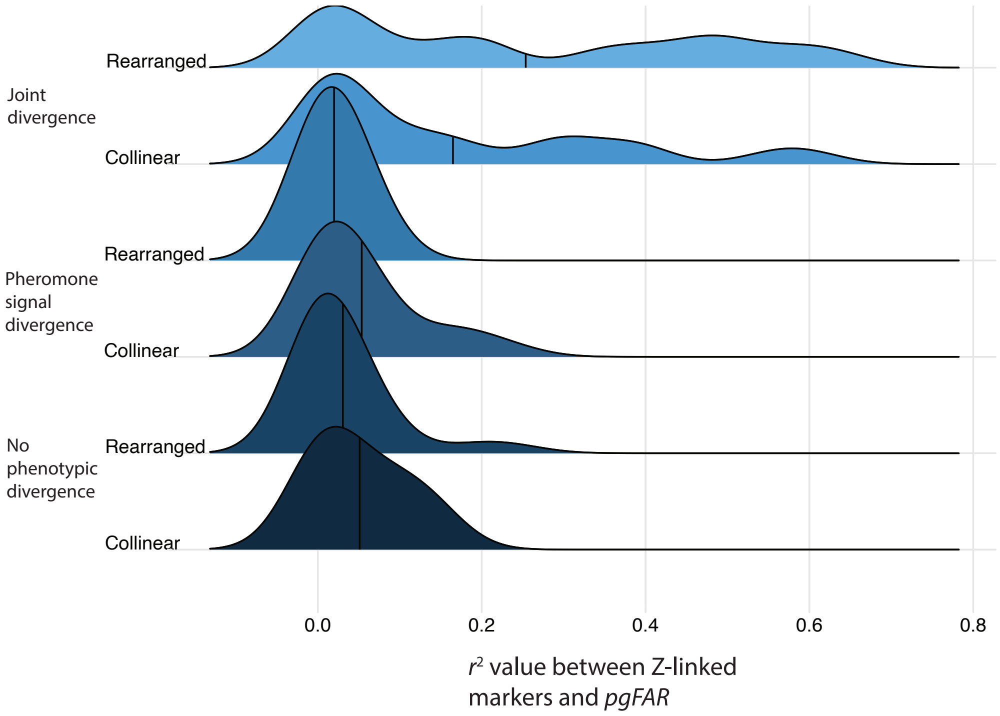


Figure S6. Linkage disequilibrium calculated within geographic populations between pgFAR and SNPs on the Z-chromosome, pooled by population type and separated by collinear vs rearranged markers. The mean is indicated by the vertical black line.

Table S1. Genome localization and other details of sequences generated by PCR primers tested and used in analysis

| Locus | Z-chromosome position Mb | Z-chromosome position cM | Features | Forward sequence | Reverse sequence |
|---|---|---|---|---|---|
| **S740_96K** | 1.165 | 4.45 | - | GATGCCTACGTCATAACATCTACCT | TAAGCCTCAGTTTTAAGTCCATTTG |
| **S196_410K** | 2.061 | 10.85 | - | TTTGTGGTTTTGGTGTTTTTATTTT | CTGTCCTGATAAACAATGTGAAATG |
| **S779_43K** | 2.929 | 10.85 | - | TCCAGCTACGTGTCAAATATCAA | AGCGTCAGTTTACAAGCTGAAAT |
| **sc216_350kb** | 3.036 | 10.85 | - | CGGTAACTAAAGGCGGAAGA | TGTCAACTTTTTGGCCACTG |
| **sc216_250kb** | 3.137 | 10.85 | - | CACAAACAAGCTGCATTTCG | AGCAGAAAATCAAACGCACA |
| **sc216_150kb** | 3.237 | 10.85 | - | TGTTCGGAGCACCATTATGA | CGCCCTACTCGAACAAAAAT |
| **sc216_100kb** | 3.287 | 10.85 | - | CATAGGGCTGTCATCCATCC | AACCAGTCATTTTCAGGGACA |
| **S390_140K** | 3.917 | 10.85 | - | CAGTCTACCCTGAAAATGGTTAGTG | TTGTATGGTGTTTGACAGATTTTTG |
| **S390_159K** | 4.155 | 10.85 | - | ACTCTACCTACTTCCATTGTTGACG | CATTACGACATGTGATTAACTTGGA |
| **S364_285K** | 4.214 | 17.59 | - | ACGCGAACTTTAAAGATTTTCTTCT | ACGTAATCAACTACACTCGACATCA |
| **sc444_225kb** | 5.269 | 25.06 | Rearrangement | CAAGACATGGTCTCCATTCG | TTTTCAATCGGGCTAAATGG |
| **sc444_250kb** | 5.294 | 25.06 | Rearrangement | ATTTGGCCCATCAAAAATTG | AGACAGTGGACGGTGTGATG |

| | | | | | |
|---|---|---|---|---|---|
| **sc444_275kb** | 5.319 | 25.06 | Rearrangement | TTAAGCA CTCGCAA TGATCG | GAACAG TAGCGA GCCACC AC |
| **sc444_300kb** | 5.344 | 25.06 | Rearrangement | GGACCTT GTTAGCT GGGAAA | CTGATG AATTCGC CGCTATT |
| **Desert** | 6.988 | 25.06 | Rearrangement | TTCAGTG CCCTCATC GTTTCTT | CCCGAA GGTGCT GTAACA AA |
| **Z11** | 6.988 | 25.06 | Rearrangement | ATCTTCCT TATTGCGT ACCA | GCATTCA GGAGCA GGAGGA |
| **CP450** | 7.231 | 25.06 | Rearrangement | AGGCCTA AAGGGAT CTTCTACC AGA | ACCCAC CATATCA ATAGCA GAGAAT |
| **REP2** | 7.231 | 25.06 | Rearrangement | AAATGGA CGATGCT GGCTGA | TTGACCG GCCAAA GTTAGG |
| **2076** | 8.076 | 25.06 | Rearrangement | GAAGCCC CAGTGTG GTGCCC | AGGGTC GCATCG AGTCCA GCT |
| **532** | 8.298 | 25.06 | Rearrangement | TGGATGT CCTCCTCT GTTTGGCT | TCACCAT CATCGG ATTCATC GGACA |
| **563** | 8.547 | 25.06 | Rearrangement | TGTTGGT AAGCGCA TCACTCA | ACATAA TTAGGG CCGTTCC ACATA |
| **P15** | 8.547 | 25.06 | Rearrangement | CGCGACC ACTCCAG TTGTTT | TGGTTTG GTCTGG AAATCTC G |
| **S645_63K** | 9.146 | 26.51 | Rearrangement | ACGAGTC TGGGTGA ATTTTATT GTA | CCAACT GAGGCT AGTTATG TAAGGA |
| **OnGH** | 9.285 | 26.51 | Rearrangement | ACGAGGG TGGCAAC GTGT | TTTCTAA GTACAG CATGTCC GTATCC |
| **P22** | 9.315 | 26.51 | Rearrangement | GATCGCG AAGCAGA | CCAAAG CGGCGG AACCAC |

| | | | | | |
|---|---|---|---|---|---|
| | | | | AGTCAGTC | |
| **S155_175K** | 10.75 | 32.5 | Rearrangement | TGATTGATGACTGATTTGGTGAT | TTGTGAATACCGGCGTAAATATC |
| **710** | 11.880 | 39.27 | Rearrangement | GCGTGCGACCCAACTCTGGA | CCATGAGCTGATCAAGGCTGATCTC |
| **AcylB** | 12.130 | 39.27 | Rearrangement | AAACTCCCGTCATCGCCG | TCTGGCCGTTCAGGATCCATTCG |
| **S1040_OR6a** | 12.240 | 40.72 | Rearrangement | TTACCATTCCATACCACCATCAT | TGTATTCCACAAGGACAAACCTC |
| **S275_OR3a** | 12.240 | 40.72 | Rearrangement | ATTATGTCCGCCGTGTAAGTAAA | GCCAAGAACGTTGATAGAATCAT |
| **S275_OR3b** | 12.240 | 40.72 | Rearrangement | AATGCATTTTGCACATAAAGGTC | TAGAACCAAATCTAATCCGCACA |
| **P25** | 12.240 | 40.72 | Rearrangement | TACGCGGGTAAGTACAGGGA | GTCGAGCGCGGCACTCATAC |
| **period** | 13.250 | 40.72 | Rearrangement | AAGAACGTCAGCGATGAAGACGGA | CAGGTAGGGCACCGACTCAGGG |
| **12295** | 14.580 | 40.72 | Rearrangement | CTGCTGATGATGAGTGATTTCCGGT | TGGTGGAGAACCATACACACTGGG |
| **12292** | 14.580 | 40.72 | Rearrangement | GATGAGCATGCGCAGACGCG | AGACGTGGTCCGACCGATCA |
| **S1038_108K** | 15.570 | 58.22 | Rearrangement | AGACGTGGTCCGACCGATCA | ATTATGCGCCGATATTTAATAACAA |
| **S48_500kb** | 16.010 | 59.87 | Rearrangement | GGTTTCCACCAAGACACCAT | CATCTTCGACACA |

| | | | | | |
|---|---|---|---|---|---|
| | | | | | GAACAGCA |
| **S48_550kb** | 16.010 | 59.87 | Rearrangement | CGGGTGTCAAACATTGGTTA | TCTTGGCCCAAATTTGTATGT |
| **S48_600kb** | 16.010 | 59.87 | Rearrangement | TGGGCAGAAAATGATGATGA | CATATCGTCGTGGTGGTTTG |
| **S48_850kb** | 16.010 | 59.87 | Rearrangement | CCGCCGGAAGTACTTAATAGC | TCCCAAACACGTGAGAGTTG |
| **LDH2** | 17.06 | 70.87 | - | CGATACCTTCTGTCCGAGAAAC | CATGGGTCTCCTTCCAGTTC |
| **S137_112K** | 17.500 | 62.22 | - | TTTTGTTTAATCCGTTTTTCACC | ACCTCAAGGTCATGGGGTATCT |
| **S137_112K** | 17.500 | 62.22 | - | GGCTTTCCCAACACAGTAATTC | TGTCGCTAAGAGAAAAAGCACA |
| **S137_112K** | 17.500 | 62.22 | - | TTGGGTGAATGCTTTTGATGTA | GGCAGACAAAATCACAAACAAG |
| **S87_90K** | 17.760 | 62.22 | - | TTGAGTTTTGGTTAAGGCCAAT | TGGGTTACCAGGGTACCATAAA |
| **S87_98K** | 17.760 | 62.22 | - | TTACGTGGTTCATACTGGAACG | ACGCAAATTAAAAGCGAATTGT |
| **S87_100K** | 17.760 | 62.22 | - | TTCAACCGAAACCATTATCAGA | AACGTTCTTGCTGTGTTGAAAA |
| **S87_102K** | 17.760 | 62.22 | - | TGTTTTCGTCTTTGTTTGTCGT | TTCCGTCTTACGTATTCCACAA |
| **S87_163K** | 17.760 | 62.22 | - | AACAGGGCACTATTAATAACGAAA | GGGATTTCAGAAAGGAAGCATA |

| | | | | | |
|---|---|---|---|---|---|
| **S50_200kb** | 18.270 | 70.87 | *bab* (Pheromone response locus) | ATCATTGGTTCCCAAAGTCG | AGGGGGCGTATCTCATTGTT |
| **S50_450kb** | 18.270 | 70.87 | *bab* | TTCCGTGATTGCAAAAACAA | CCGGAGTCACTTTGGCTAGT |
| **S50_550kb** | 18.270 | 70.87 | *bab* | GACGGAACCCGTTGTCTTAG | TCGTTACGATGCCAGTACGA |
| **S662_45K** | 18.880 | 70.87 | *bab* | GCATTTGAAATCTCACCTTGCTA | CTGCTCCCTCCTGGAAATCTA |
| **S662_149K** | 18.880 | 70.87 | *bab* | TTAGACATCCAAACTTTCGCATT | TCCGCTAAATACTATTGG |
| **S229_191K** | 19.260 | 74.6 | - | AAATACCGAAAGGTCCTAACATACC | AGGCTCGCGTAAGTATCATTT |
| **S229_199K** | 19.260 | 74.6 | - | GCTTTGGTATCATTATGTCCTTTGT | CCTCTCGAGTTCTTTTTCAC |
| **EIPB** | 20.190 | 82.7 | - | GTACAGCAGGTCGTGGAGTT | CCATCATCGAAGACGAGGC |
| **c00710** | 20.470 | 82.7 | - | CTAGCTCGGTTTCACATTTCAAGT | TACCCACCGTCCAACACATAAG |
| **647** | Likely autosomal | - | - | TGCAAGAAGTGGTCCAGGCACT | GCAAGATGCCGAAGTCGCTC |
| ***pgFAR*** | Autosomal | - | Pheromone production gene | GGTGGCATGGGGACGGTACAG | TTTGGATTAAATTAATTTTAAAC |
| **sc178_200kb** | Autosomal | - | Pheromone production scaffold | ATGAGATGTGGTTGCACGAA | TCATTGTTTCAGCCACTGGA |
| **sc178_300kb** | Autosomal | - | Pheromone production scaffold | ACAAGTTATGCCGGACAAGG | CTTACAGATGCTGGCAATCG |

| | | | | | |
|---|---|---|---|---|---|
| **sc178_400kb** | Autosomal | - | Pheromone production scaffold | CGTCTGCGAGTGTCCATTTA | CCCAATCCCAACGAAAATAA |
| **sc178_500kb** | Autosomal | - | Pheromone production scaffold | TGATGATGAATCGCTTTTGC | CCGTTTCGAACCACTACACC |
| **sc178_550kb** | Autosomal | - | Pheromone production scaffold | TAGCTGCTGGAGGATTTTGG | TGGATGAGAAAGCGGAAGAT |
| **cry2** | Autosomal | - | - | CGGTGCGGCCGTCGATCCATC | AACCAGTCATTTTCAGG |
| **G26A** | Autosomal | - | - | AGATCTCGCACGAATTTTGAATG | ACGCCTATTATTACGTCAGG |

Table S2. STRUCTURE output statistics based on the Evanno method (Evanno et al. 2005).

| K | Replicates | Mean LnP(K) | Standard deviation LnP(K) | Ln'(K) | \|Ln''(K)\| | Delta K |
|---|---|---|---|---|---|---|
| 1 | 50 | -57459.252 | 6.8696 | NA | NA | NA |
| 2 | 50 | -49713.204 | 4.7239 | 7746.048 | 6152.868 | 1302.50176 |
| 3 | 50 | -48120.024 | 9.548 | 1593.18 | 2566.848 | 268.836187 |
| 4 | 50 | -49093.692 | 1586.3296 | -973.668 | 339.462 | 0.213992 |
| 5 | 50 | -49727.898 | 2741.6036 | -634.206 | 53.446 | 0.019494 |
| 6 | 50 | -50308.658 | 1384.6757 | -580.76 | 259.016 | 0.187059 |
| 7 | 50 | -50630.402 | 1451.1248 | -321.744 | NA | NA |

The grey bar indicates the most likely number of inferred clusters.
